# E-cadherin in developing murine T cells controls spindle alignment and differentiation during β-selection

**DOI:** 10.1101/2022.06.14.496211

**Authors:** Mirren Charnley, Amr H Allam, Lucas M Newton, Patrick O Humbert, Sarah M Russell

**Author notes:** now at Olivia Newton-John Cancer Research Institute and La Trobe University School of Cancer Medicine, Heidelberg, VIC, 3084, Australia.

## Abstract

A critical stage of T cell development is β-selection; at this stage the TCRβ chain is generated and the developing T cell starts to acquire antigenic specificity. Progression through β-selection is assisted by a low affinity interaction between the nascent TCRβ chain and peptide presented on stromal MHC and external cues provided by the niche, including Notch and CXCR4. In this study, we reveal the importance of a new cue within the murine developing T cell niche which is critical for T cell development. E-cadherin mediates cell-cell interactions and influences cell fate in many developmental systems. In developing T cells E-cadherin contributed to the formation of an immunological synapse and the alignment of the mitotic spindle with the polarity axis during division, which facilitated subsequent T cell development. Collectively, these data highlight a new aspect of the developing T cell niche and provide insights into the role of E-cadherin in the β-selection stage of T cell development.

## Introduction

T cell development requires both the cell fate specification process common to all cells in a multicellular organism, and the creation of a unique, genomically recombined T cell receptor (TCR) (Shah and Zuniga-Pflucker 2014). The process of genomic recombination means that errors in this TCR diversification can lead to leukemia, immunodeficiencies or autoimmunity (Michie and Zúñiga-Pflücker 2002, Chann and Russell 2019). To prevent such errors, T cell development occurs in a series of highly orchestrated stages, in different regions of the thymus. The first such genomic recombination occurs at the β-selection stage, when the TCRβ chain is created. This stage is characterised by major and abrupt changes in cell phenotype and fate and includes a β-selection checkpoint which many cells fail to pass (Dutta, Zhao et al. 2021). Passage through the β-selection checkpoint is controlled by several external cues comprising a restricted niche in the thymus (Ciofani and Zuniga-Pflucker 2005, Janas, Varano et al. 2010, Charnley, Ludford-Menting et al. 2020, Allam, Charnley et al. 2021). Competition for this niche influences cell fate and protects from leukemia (Boehm 2012, Martins, Busch et al. 2014, Paiva, Sousa et al. 2021), but the composition and function of cues provided by the niche are not well understood.

It has recently become clear that, contrary to previous dogma, β-selection involves a low affinity interaction between the nascent TCRβ chain and peptide presented on stromal MHC (Mallis, Bai et al. 2015). This interaction can induce the formation of an immunological synapse, similar to that of mature T cells, for passage through the β-selection checkpoint (Allam, Charnley et al. 2021). Other receptors such as Notch and CXCR4 can facilitate pre-TCR signalling and the polarisation of the developing T cell towards the stromal cell bearing MHC-peptide during β-selection (Pham, Shimoni et al. 2015, Charnley, Ludford-Menting et al. 2020, Allam, Charnley et al. 2021). Together, these signals coordinate polarity to enable asymmetric cell division of the developing T cell during β-selection, again supporting the notion of a niche that controls cell fate (Pham, Shimoni et al. 2015, Charnley, Ludford-Menting et al. 2020). However, given the lack of coreceptors at the β-selection stage and the weak binding of the pre-TCR compared with the TCRαβ of mature T cells, it seems likely these cells would need more help to maintain an interaction sufficiently strong to support TCR signalling and polarisation, particularly in the context of competition for the niche.

We therefore explored whether other cell attachment receptors might enhance the adhesion of developing T cells in the thymic niche during β-selection. VCAM1 has the capacity to provide such an interaction, being presented by cortical stromal cells and binding the α4 integrin on early T cell progenitors (Prockop, Palencia et al. 2002). Indeed, the VCAM1-α4 integrin interaction is critical for cells just prior to the β-selection stage (termed DN2) to migrate to the outer cortex where they commence β-selection (Prockop, Palencia et al. 2002), and synergizes with Notch ligands for *in vitro* differentiation to the DN3 stage (Shukla, Langley et al. 2017, Michaels, Edgar et al. 2021). However, expression of the α4 integrin drops sharply from the DN2 stage (Michaels, Edgar et al. 2021), and polarisation to DL4 is inhibited, rather than enhanced, by VCAM1 in DN3a undergoing asymmetric cell division (Charnley, Ludford-Menting et al. 2020).

A second candidate to facilitate an interaction with stromal cells is E-cadherin. E-cadherin is a well-known mediator of cell–cell interactions through its extracellular domain (Halbleib and Nelson 2006). Furthermore, it acts as an external cue to dictate intracellular polarisation and control mitotic spindle orientation during division (den Elzen, Buttery et al. 2009, Desai, Gao et al. 2009, Dupin, Camand et al. 2009, Charnley, Kroschewski et al. 2012, Shahbazi and Perez-Moreno 2015). It is expressed by thymic epithelial cells and developing T cells (Lee, Sharrow et al. 1994, Muller, Luedecker et al. 1997, de Yzaguirre, Hernandez et al. 2006). Treatment of disaggregated thymi with an antibody that blocked homotypic E-cadherin interactions (but not E-cadherin-α_E_β_7_ interactions) impaired the structure of the thymus, and inhibited the ability of thymic epithelial cells to promote differentiation to the DP stage of T cell development, although it was not clear which precursor stage was blocked (Lee, Sharrow et al. 1994, Muller, Luedecker et al. 1997). In further support for a possible role in β-selection, two functional partners of E-cadherin that influence progression through β-selection and T cell development, Scribble and β-catenin (Pike, Kulkarni et al. 2011, Ma, Wang et al. 2012, Parsons, Patel et al. 2019), also coordinate polarity with spindle orientation in many cell types (Shahbazi and Perez-Moreno 2015, Bonello and Peifer 2019). Together, these observations suggested that E-cadherin might facilitate adhesion and polarisation to stromal cells to enable passage through the β-selection checkpoint.

Here we demonstrate that E-cadherin alters T cell development at β-selection by coordinating adhesion, polarity and spindle orientation during asymmetric cell division. We characterised the role of E-cadherin during T cell development using the OP9 stromal cell line expressing Notch ligand, DLL1 (Schmitt and Zuniga-Pflucker 2002, Schmitt and Zuniga-Pflucker 2006, Dervovic, Ciofani et al. 2012), combined with an inducible Cre recombinase based knockout system to prevent E-cadherin signalling, and surfaces functionalised with E-cadherin. We show that E-cadherin was critical for the regulation of mitotic spindle orientation in dividing T cells and coordinating it with polarity of proteins during ACD. Inhibiting E-cadherin hindered progression through the β-selection checkpoint and subsequently reduced T cell differentiation. This highlights a new aspect of microenvironment control during T cell development and provides a means by which the low affinity interactions of pre-TCR with MHC-peptide might be sustained in the absence of coreceptors.

## Results

### Exogenous E-cadherin can induce polarisation of DN3 cell fate determinants via homotypic adhesions

To determine whether E-cadherin might contribute to cell polarity during β-selection, we first assessed whether exogenous E-cadherin supported polarity (**Fig. 1A**). DN3a cells were cultured on E-cadherin surfaces for 2 or 15 hours and rinsed to remove any non-adherent cells. The DN3a cells attached to the surfaces within 2 hours (**Fig. S1A**). DN3a cells polarised their microtubule organising centre (MTOC) to the cell-surface interface in approximately 60% of cells (**Fig. 1B, C**), which was comparable to the level of MTOC polarisation previously observed on DL1 functionalised surfaces (Charnley, Ludford-Menting et al. 2020). We have shown that DN3a cells polarise cell fate determinants during cell division in response to specific cues from the stromal cells, and this can be mimicked in the case of Notch ligands by surfaces functionalised with Fc-DL1 (Pham, Shimoni et al. 2015, Charnley, Ludford-Menting et al. 2020). Similarly, DN3a cells on E-cadherin functionalised surfaces showed polarisation of endogenous fate determinants, Numb and α-adaptin, but not the control marker, CD25, during interphase (Fig 1B, C) and division **(Fig. 1D)**. Polarisation was also observed using time lapse analysis of dividing DN3a cells expressing cherry-tagged Numb or α-adaptin cultured on E-cadherin functionalised surfaces (**Fig. 1E and S1B-E**). Polarisation in response to E-cadherin was comparable to polarisation in response to OP9-DL1 stromal cells (Charnley, Ludford-Menting et al. 2020) or cell contacts with neighbouring DN3a cells, but not on PA coated surfaces (Fig 1E). Hence, E-cadherin is sufficient to trigger polarisation during division.

**Figure 1.**
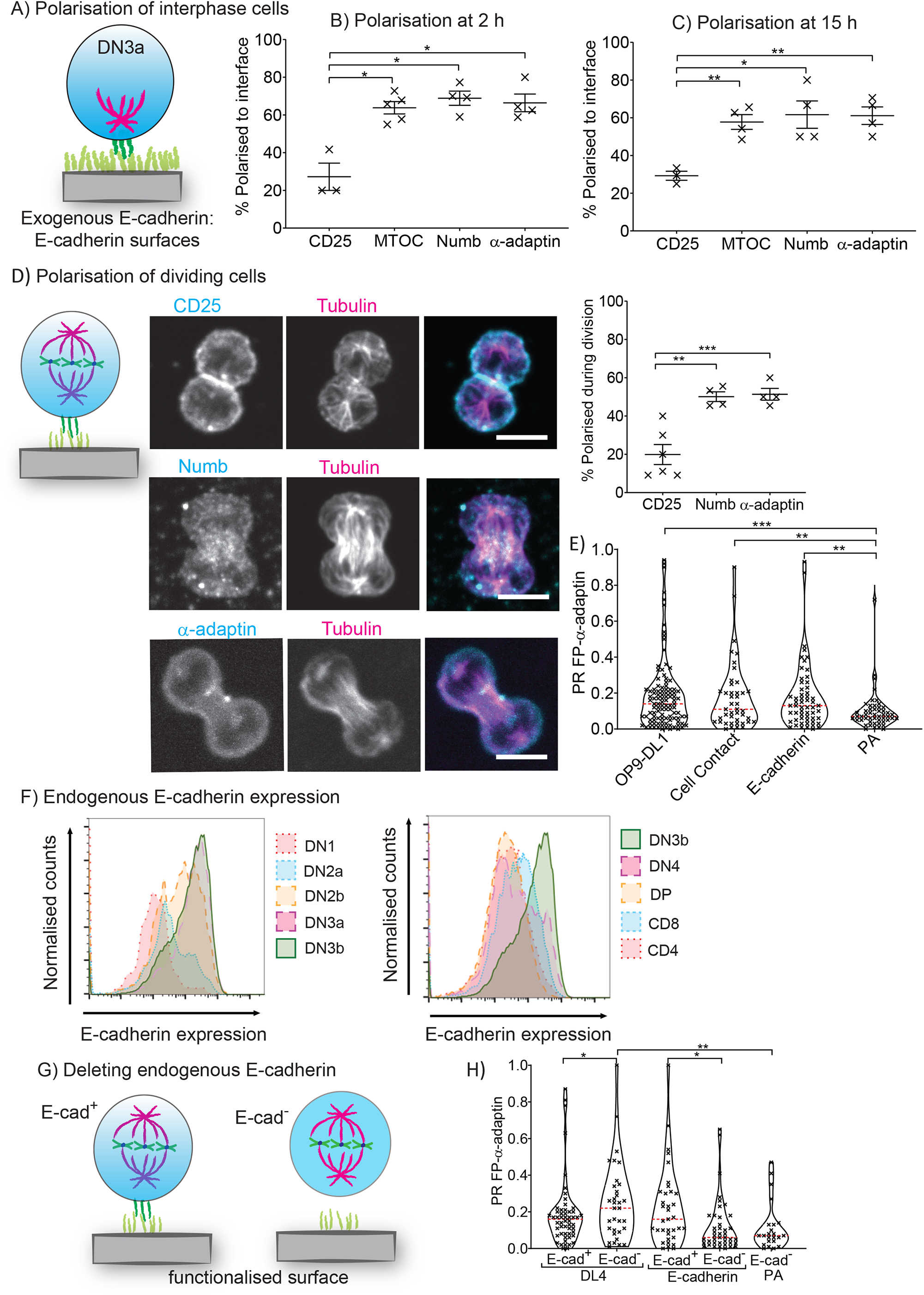
Exogenous E-cadherin can mediate polarisation of fate determinants via homotypic interactions. To determine whether E-cadherin could support adhesion and polarisation of developing T cells DN3a cells were re-seeded onto PA + E-cadherin coated surfaces and fixed and stained after 2 or 15 hours or imaged for 20 hours using time lapse microscopy. (A) Schematic of the experimental set up used to test the role of exogenous E-cadherin in polarisation. Approximately half of the cells polarised the MTOC (identified by tubulin staining) to the cell-substrate interface after (B) 2 and (C) 15 hours contact with the surfaces. Numb and α-adaptin, but not CD25, were polarised in a high proportion of cells during interphase. (D) Representative projected z stack images of dividing DN3a cells with corresponding quantification on the right, bar = 10 μm. Numb and α-adaptin were polarised in approximately 50% of dividing cells. Number of cells analysed: interphase: CD25 = 44 (2 h), 38 (15 h), α-adaptin = 65 (2 h), 53 (15 h) and Numb = 84 (2 h), 55 (15 h); divisions: CD25 = 61, α-adaptin = 41 and Numb = 48. (E) Violin plot of dividing DN3a cells. The distribution of PR values was greater in the presence of E-cadherin and was comparable to polarisation in response to OP9-DL1 stromal cells (Charnley, Ludford-Menting et al. 2020). (F) E-cadherin expression was assessed at the different stages of T cell development was using flow cytometry. E-cadherin increased as the developing T cell progressed from to DN2b, peaked at DN3 and started to decrease again at DN4 onwards. (G) Schematic of the experimental set up used to test the role of endogenous E-cadherin in polarisation. (H) DN3a cells were transduced with α-adaptin either GFP (E-cad^+^) or GFP-Cre (E-cad^-^) and seeded onto PA, PA + DL4 or PA + E-cadherin. E-cadherin knockout reduced the level of ACD on E-cadherin coated surfaces. Flow cytometry plots of one representative experiment are shown, n = 3-6 independent experiments. Magenta line on violin plots indicates median value. *p < 0.05, ** p < 0.01, ***p < 0.001 (t test or KS test for violin plots).

This effect of exogenously presented E-cadherin on polarisation might be mediated either by homotypic (E-cadherin-E-cadherin) interactions, or heterotypic interactions such as between E-cadherin and integrins. To assess this, we first explored the kinetics of expression of E-cadherin in developing T cells using flow cytometry. E-cadherin expression was low in DN1 and DN2a cells; peaked at DN3 and declined from DN4 (**Fig. 1F**). This expression pattern is consistent with E-cadherin being integral to the β-selection checkpoint. An alternative binding partner for E-cadherin is the integrin αEβ7. β7 was highly expressed at DN2b but dropped as the cells progressed to DN3 and DN4 and αE expression was consistently low (**Fig. S1F**). This suggests that the polarisation observed was mediated by homotypic E-cadherin-E-cadherin interactions. To confirm this, we knocked out E-cadherin using an inducible Cre recombinase system (**Fig. 1G**). Initially, we confirmed that transduction with GFP-Cre correlated with an abrogation of E-cadherin expression in DN3 cells, and the expression of E-cadherin was hindered up to 11 days after sorting (**Fig. S2A**). Developing T cells were transduced with cherry α-adaptin and either GFP (E-cadherin^+^) or GFP-Cre (E-cadherin^-^), sorted for DN3a cells and re-seeded onto OP9-DL1 stromal cells for time lapse imaging (**Fig. 1H and S2B-F**). In dividing DN3a cells, α-adaptin was polarised in response to E-cadherin (as in Fig 1E) and to DL4 (as previously found (Charnley, Ludford-Menting et al. 2020)). Deletion of E-cadherin in the DN3a cells prevented polarisation in response to exogenous E-cadherin but did not prevent polarisation in response to DL4. Together these data indicate developing T cells express E-cadherin during β-selection and suggest that homotypic E-cadherin interactions can induce polarisation of cells undergoing β-selection.

### Endogenous E-cadherin is polarised towards OP9-DL1 cells and promotes immunological synapse stability but not polarisation of fate determinants during cell division

Having shown that homotypic interactions can promote polarisation during β-selection, we next assessed the role of endogenous E-cadherin in the DN3a response to OP9-DL1 stromal cells, which also express E-cadherin (**Fig. S3A)**. Endogenous E-cadherin was polarised to the cell-cell interface in DN3a cells cultured on OP9-DL1 stromal cells in interphase (**Fig. 2A**) and during division (**Fig. 2B**). Thus, E-cadherin is polarised to the cell contact with the OP9-DL1 stromal cell.

**Figure 2.**
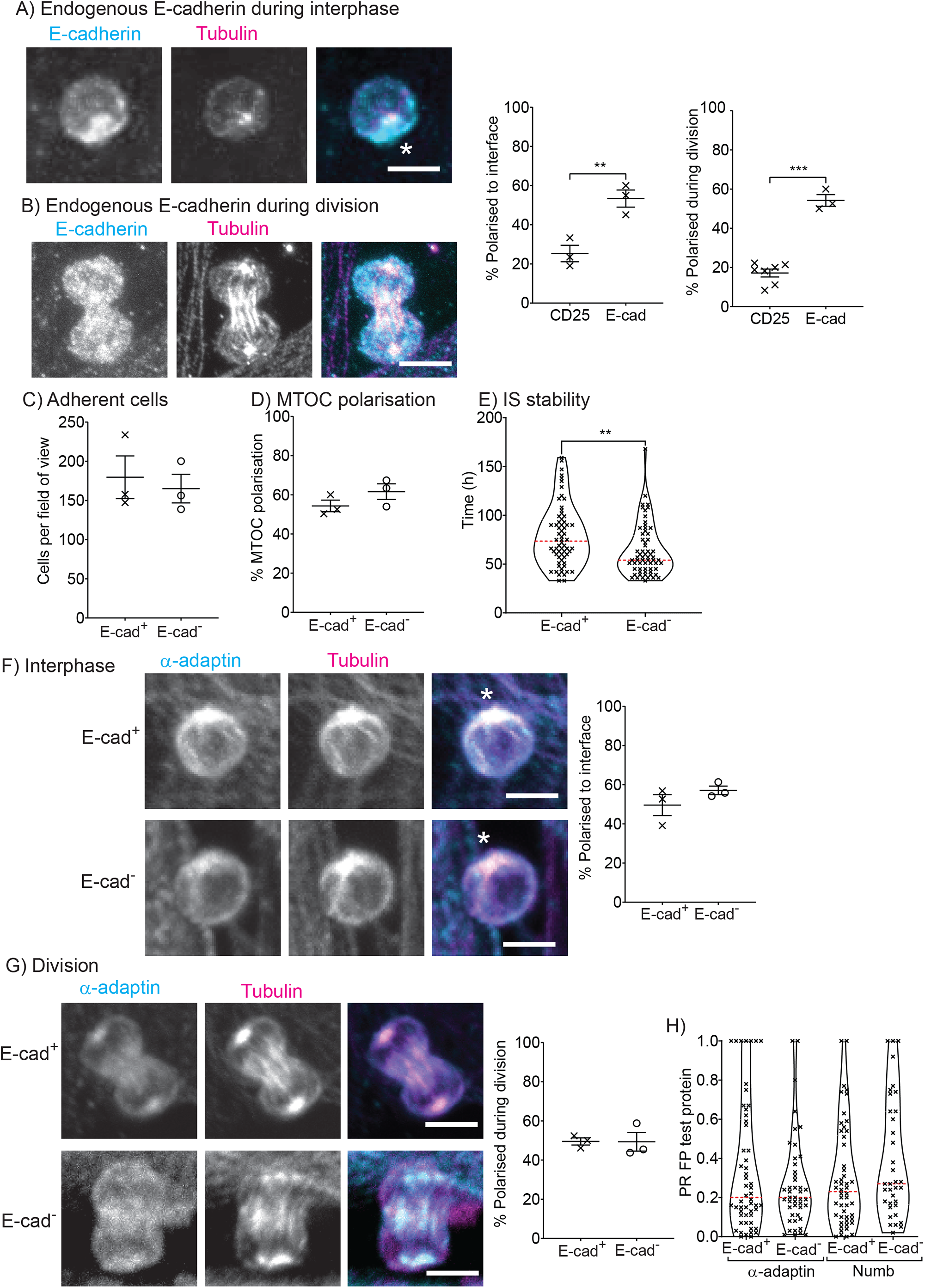
Endogenous E-cadherin in DN3A cells is polarised towards OP9-DL1 cells and required for immunological synapse stability but not for polarisation of fate determinants during β-selection. DN3a cells were seeded onto OP9-DL1 stromal cells and analysed for the spatial distribution of endogenous E-cadherin. (A) Endogenous E-cadherin was polarised to the cell-cell contact and (B) asymmetrically distributed between the two daughter cells in approximately 50% of dividing cells. Non-polarised control protein CD25 is shown for comparison (Charnley, Ludford-Menting et al. 2020). Number of divisions analysed: CD25 = 86 and E-cadherin = 47. To inhibit E-cadherin developing T cells from E-cadherin^flox/flox^ mice were transfected with GFP (E-cad^+^) or GFP-Cre (E-cad^-^) on Day 4 and sorted on Day 8. DN3a cells were re-seeded onto OP9-DL1 stromal cells and after 15 h fixed and stained for tubulin and α-adaptin. E-cadherin knockout did not affect (C) the number of attached DN3a cells or (D) the proportion of the cells that polarised the MTOC to the cell-cell interface. (E) To determine the role of E-cadherin in immunological synapse stability DN3a cells transfected with cherry-tubulin were imaged using time lapse microscopy (**see video 1**). Cells were triaged for cells in which MTOC was recruited to the interface with an OP9-DL1 cell for at least 30 minutes and the cells were tracked over time to determine the time for which the MTOC remained at the synapse. The duration of the immunological synapse was reduced in the absence of E-cadherin. (F) α-adaptin was polarised during interphase and (G) in approximately 50% of dividing cells and this polarity was not affected by inhibiting E-cadherin. Number of divisions analysed: 48 (α-adaptin, E-cad^+^), 44 (α-adaptin, E-cad^-^). (H) DN3a cells were transduced with α-adaptin or Numb and either GFP or GFP-Cre and imaged for 20 hours using time lapse microscopy. The distribution of PR values was similar regardless of whether the cells were transduced with GFP or GFP-Cre. Magenta line on violin plots indicates median value. Representative images of DN3a cells cultured on OP9-DL1 is shown on the left with corresponding quantification on the right, *indicates the interface between the DN3a and OP9-DL1 stromal cells, bar = 10 μm. n = 3 -5 independent experiments, ** p < 0.01,***p < 0.001.

The number of cells that adhered to the stromal layer was not affected by the deletion of E-cadherin (**Fig. 2C**). During β-selection signalling proteins including MTOC, Notch, CXCR4 and pre-TCR, are coordinated into a signalling platform, reminiscent of the immunological synapse in mature T cells (Allam, Charnley et al. 2021). To determine whether this immunological synapse was altered in the absence of E-cadherin, we first assessed recruitment of the MTOC to the interface between the DN3a cell and stromal cell 15 hours after seeding and found no difference (**Fig. 2D**). However, time lapse imaging showed that the stability of this MTOC recruitment was slightly reduced in the absence of E-cadherin (**Fig. 2E**). Together, these findings suggest that E-cadherin is polarised and can contribute to immunological synapse stability during T cell development.

Previously, we have demonstrated that α-adaptin and Numb are polarised in dividing T cells (Pham, Shimoni et al. 2015, Charnley, Ludford-Menting et al. 2020) and exogenous E-cadherin can act as a cue to trigger this response (Fig. 1). Hence, we tested whether endogenous E-cadherin in the developing T cells was needed for polarisation of α-adaptin and Numb in response to stromal cells. As expected, wild type DN3a cells (transduced with the control plasmid, GFP) showed strong polarisation of both α-adaptin and Numb during interphase and mitosis (**Fig. 2F-H and S3B, D**). E-cadherin deficient DN3a cells showed remarkably similar levels of polarisation of both proteins during division (**Fig. 2F-H and S3C, E**). This indicates that when DN3a cells are cultured on OP9-DL1 stromal cells, the deletion of endogenous E-cadherin did not affect the polarity of Numb and α-adaptin during interphase or division. Thus, E-cadherin in the DN3a cells is recruited to the interface with the stromal cells, but not required for stromal-induced polarity of fate determinants during division.

### E-cadherin-associated regulators of the mitotic spindle are polarised towards OP9-DL1 cells, and endogenous E-cadherin is required for their polarisation

E-cadherin is known to act as a critical link between the microenvironment and spindle orientation in epithelial cells (Halbleib and Nelson 2006, Martin-Belmonte and Perez-Moreno 2011, Matsuzaki and Shitamukai 2015, Shahbazi and Perez-Moreno 2015). To decipher whether E-cadherin played a similar role in developing T cells we examined the localisation of three proteins, β-catenin, adenomatous polyposis coli (APC) and nuclear mitotic apparatus (NuMA), which have previously been demonstrated to bridge E-cadherin contacts with the cytoskeleton to control spindle orientation in epithelial cells (Olmeda, Castel et al. 2003, den Elzen, Buttery et al. 2009, Shahbazi and Perez-Moreno 2015, Kiyomitsu and Boerner 2021).

β-catenin is typically concentrated at the cell-cell contact or cortex in dividing cells, where it interact with E-cadherin and the cytoskeleton (Shahbazi and Perez-Moreno 2015). When E-cadherin was inhibited it became diffusely localised in the cytoplasm (Olmeda, Castel et al. 2003, den Elzen, Buttery et al. 2009). In developing T cells β-catenin localisation was concentrated at the cortex during division, reminiscent of its localisation in epithelial cells (**Fig. 3A, B** and **S4A**). The deletion of E-cadherin increased β-catenin in the cytoplasm (**Fig. 3B** and **S4B**). Furthermore, β-catenin was polarised in approximately 60% of dividing DN3a cells, which was reduced in the absence of E-cadherin. Thus, β-catenin is predominantly cortical in DN3 cells, and this cortical localisation is reduced by depletion of endogenous E-cadherin.

**Figure 3.**
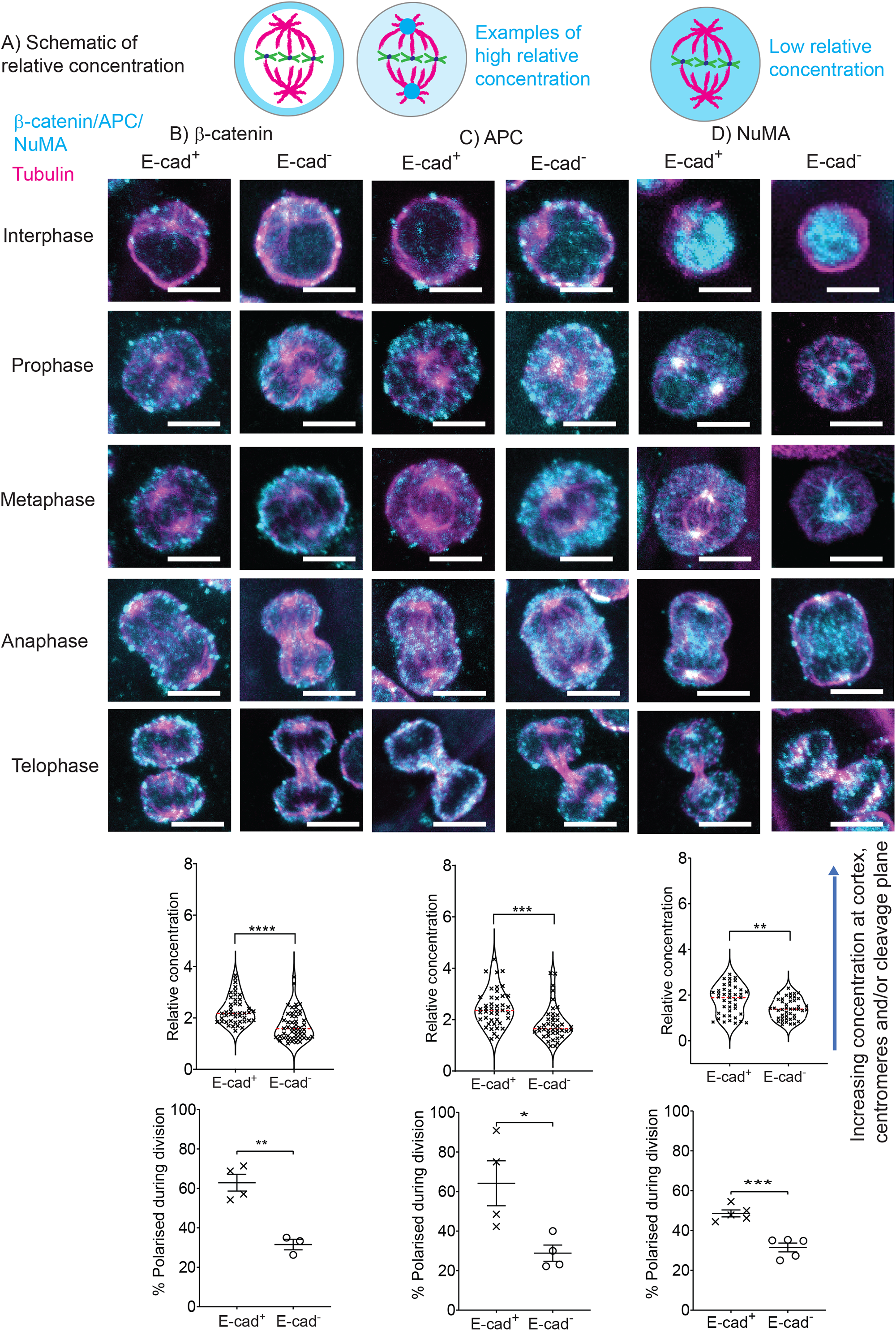
E-cadherin-associated regulators of the mitotic spindle are polarised towards OP9-DL1 cells, and endogenous E-cadherin is required for their polarisation. Developing T cells from E-cadherin^flox/flox^ mice were transfected with GFP (E-cad^+^) or GFP-Cre (E-cad^-^) on Day 4 and sorted on Day 8. DN3a cells were re-seeded onto OP9-DL1 stromal cells and after 15 h fixed and stained for tubulin and either β-catenin, adenomatous polyposis coli (APC) or NuMA. (A) The fluorescent intensity of the protein of interest was measured at the cortex, midbody and centromeres and ratioed to the fluorescent intensity of the cytoplasm. Higher scores indicate that the protein was concentrated in these regions, while lower scores indicates the protein was diffuse. Schematic of high relative concentration where the protein of interest is concentrated at the cortex or centromeres versus low relative concentration where the protein is diffuse through the dividing cell. Inhibiting E-cadherin correlated with a more diffuse distribution of (A) β-catenin, (B) adenomatous polyposis coli (APC) or (C) NuMA. Inhibiting E-cadherin also reduced the proportion of dividing cells that polarised all three spindle proteins. Number of divisions analysed for polarisation: 74 (APC, E-cad^+^), 68 (APC, E-cad^-^), 54 (β-catenin, E-cad^+^) 70 (β-catenin, E-cad^-^), 116 (NuMA, E-cad^+^) and 86 (NuMA, E-cad^-^). Representative images of the spindle regulators in DN3a cells cultured on OP9-DL1 is shown with corresponding quantification is shown below bar = 10 μm. n = 3 -5 independent experiments; magenta line on violin plots indicates median value. *p < 0.05, ** p < 0.01, ***p < 0.001, ****p < 0.0001 (t test or KS test for violin plots).

In epithelial cells adenomatous polyposis coli (APC) binds to β-catenin and microtubule binding proteins, such as EB1, and during division it concentrates at the cell cortex or at the centromeres (Olmeda, Castel et al. 2003, den Elzen, Buttery et al. 2009). The C. elegans homologue of APC, APR-1 localises to the anterior cortex of the dividing zygote to position the mitotic spindle (Sugioka, Fielmich et al. 2018). In developing T cells, APC was asymmetric and at the cortex during division but was not observed at the centromeres (**Fig. 3C** and **S5A**). Similar to previous research in epithelial cells (den Elzen, Buttery et al. 2009), the absence of E-cadherin caused a reduction in the level of APC at the cortex relative to the cytoplasm and reduced its polarisation in dividing DN3a cells (**Fig. 3C** and **S5B**). Thus, endogenous E-cadherin is also required for the correct spatial localisation of APC.

NuMA forms a complex with LGN and Gα at the E-cadherin contact to direct the position of the mitotic spindle (Gloerich, Bianchini et al. 2017, Kiyomitsu and Boerner 2021). In epithelial cells NuMA is concentrated in the nucleus during interphase and translocates to the centromeres and cortex during division (den Elzen, Buttery et al. 2009, Poulson and Lechler 2010, Gloerich, Bianchini et al. 2017, Kiyomitsu and Boerner 2021). NuMA was strongly recruited to the spindle poles in developing T cells, suggesting that NuMA might also play a role in spindle orientation during T cell development (**Fig. 3D** and **S6A, B**). Similar to the other E-cad-associated proteins, and in contrast to epithelial cells (den Elzen, Buttery et al. 2009), E-cadherin was required for optimal concentration of NuMA at the centromeres (Fig. 3D and S6A, B). NuMA was also polarised distributed across the two daughter cells in an E-cadherin dependent manner.

Taken together, the distribution of these three regulators of the mitotic spindle during division was reminiscent of their distribution in epithelial cells, and endogenous E-cadherin is required for the correct distribution of these proteins. The polarisation of β-catenin, APC and NuMA during cell division suggested that they might play a role in linking polarity to spindle orientation during asymmetric cell division, as has been observed in *Drosophila* and *C. elegans* (Bowman, Neumuller et al. 2006, Mizumoto and Sawa 2007). The dependence of their polarisation on E-cadherin expression was in contrast to the lack of dependence on E-cadherin of the fate determinants, Numb and α-Adaptin (Fig. 2). these findings raise the possibility that E-cadherin might act early in division to coordinate the axis of polarity with the orientation of the mitotic spindle during β-selection.

### E-cadherin is required for alignment of the mitotic spindle with the polarity axis during division

To assess whether E-cadherin influences orientation of the mitotic spindle in developing T cells, we measured the angle of division relative to the polarity cue in dividing DN3a cells cultured on OP9-DL1 stromal cells (**Fig. 4A**). As previously observed (Pham, Shimoni et al. 2015, Charnley, Ludford-Menting et al. 2020), higher PR values correlated with higher spindle angles in the presence of E-cadherin (**Fig. 4B**). However, when E-cadherin was deleted this correlation was lost (**Fig. 4C**). The loss of E-cadherin resulted in fewer divisions perpendicular to the cue (division angles greater than 45°, 37% for E-cad^-^ *versus* 46% for E-cad^+^), and in the cells that did divide perpendicular this did not correlate with a high PR, in contrast to the E-cad^+^ control (**Fig. 4D**). As has been previously observed in HeLa cells (Charnley, Anderegg et al. 2013), spindle misorientation correlated with a slight increase in the time that the cell took to progress through division (**Fig. 4E**). Thus, E-cadherin was critical for the correct orientation of the mitotic spindle. Taken together these results indicate that E-cadherin is required for the orientation of the spindle, perhaps by altering the activity of spindle regulators, β-catenin, APC and NuMA. Consequently, the coordination of polarity of cell fate proteins relative to the mitotic spindle was disrupted in the absence of E-cadherin.

**Figure 4.**
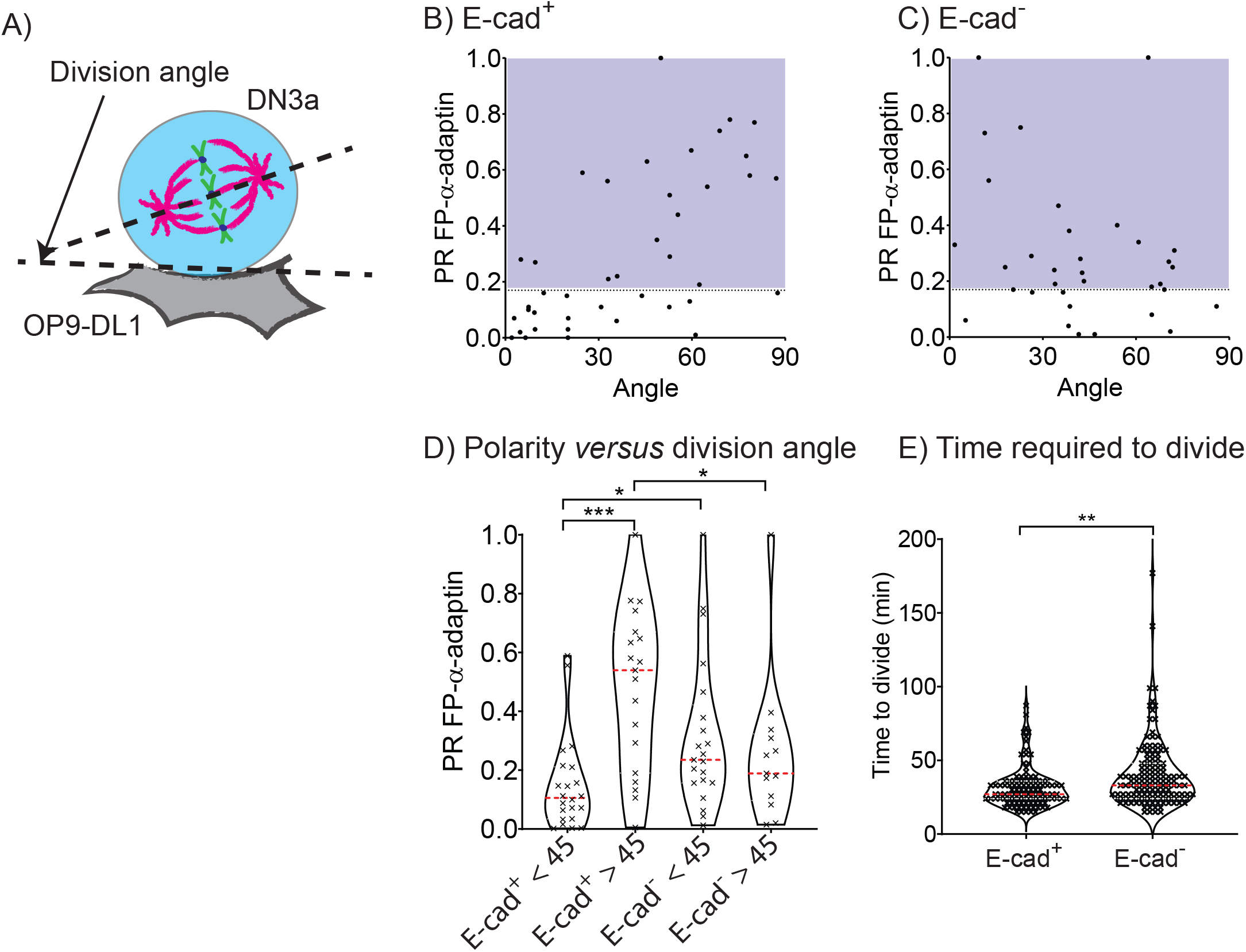
E-cadherin is required for alignment of the mitotic spindle with the axis of polarity during division. Developing T cells from E-cadherin^flox/flox^ mice were transfected with GFP or GFP-Cre and cherry α-adaptin on Day 4 and sorted on Day 8. DN3a cells were re-seeded onto OP9-DL1 stromal cells and imaged for 20 hours using time lapse microscopy. (A) To determine the effect on spindle orientation the division angle, defined as the angle between the DN3a cell and OP9-DL1 stromal cell interface and the long axis of the dividing cell, was measured and plotted *versus* PR. The blue shaded region indicates divisions that were assigned as polarised when the 0.17 cut-off was applied. (B) Higher divisions angles correlated with higher PRs in cells transduced with (B) GFP (Ecad^+^) but not (C) GFP-Cre (E-cad^-^). (D) The same trend was observed when we compared the spread in PR values for cells which divided with a division angle < 45° *versus* > 45°. (E) Deleting E-cadherin also slightly increased the time required for the cells to divide (**see video 2**). n = 2 - 5 independent experiments, magenta line on violin plots indicates median value; *p < 0.05, ** p < 0.01, ***p < 0.001 (KS test).

### E-cadherin is required for T cell development

Deleting E-cadherin slightly altered the duration of the immunological synapse and hindered the correct alignment of the mitotic spindle with the axis of polarity for fate regulators, which we postulated would have a downstream effect on T cell development. To test this *in vitro* we used the Cre recombinase E-cadherin knockout system and OP9 coculture to support T cell development. On Day 8 of the coculture the developing T cells were sorted for GFP+ cells (reflecting expression of Cre or control GFP) and re-seeded on OP9-DL1 stromal cells and analysed for differentiation using flow cytometry. Data from 1 representative experiment is shown in **Fig. S7**. To compare across experiments we derived ratios for the E-cad^-^ (GFP-Cre) versus the E-cad^+^ (GFP-control transduced) cells from the floxed (E-cad^FL^) mice for each culture, where an increase in the ratio indicates an increase in E-cadherin-deficient cells (E-cad^FL:Cre^ compared with E-cad^FL:WT^) (**Fig. 5 to 7**). To control for the well-known Cre recombinase toxicity (Schmidt-Supprian and Rajewsky 2007, Carow, Gao et al. 2016) the results were compared to developing T cells from E-cad^WT^ mice (not floxed) transduced with either GFP or GFP-Cre (E-cad^WT:WT^ and E-cad^WT:Cre^). The toxicity of the Cre recombinase is evident in the wild type control; the Cre/WT ratio was less than 1 indicating the cell number and differentiation was reduced after transduction with Cre recombinase. However, this toxicity effect was not stage specific and by comparing between the Cre/WT ratios for the wild type (E-cad^WT^) and E-cadherin KO (E-cad^FL^) it was possible to observe clear effects of the E-cadherin KO beyond those ascribable to Cre recombinase toxicity (**Fig. 5A)**.

**Figure 5.**
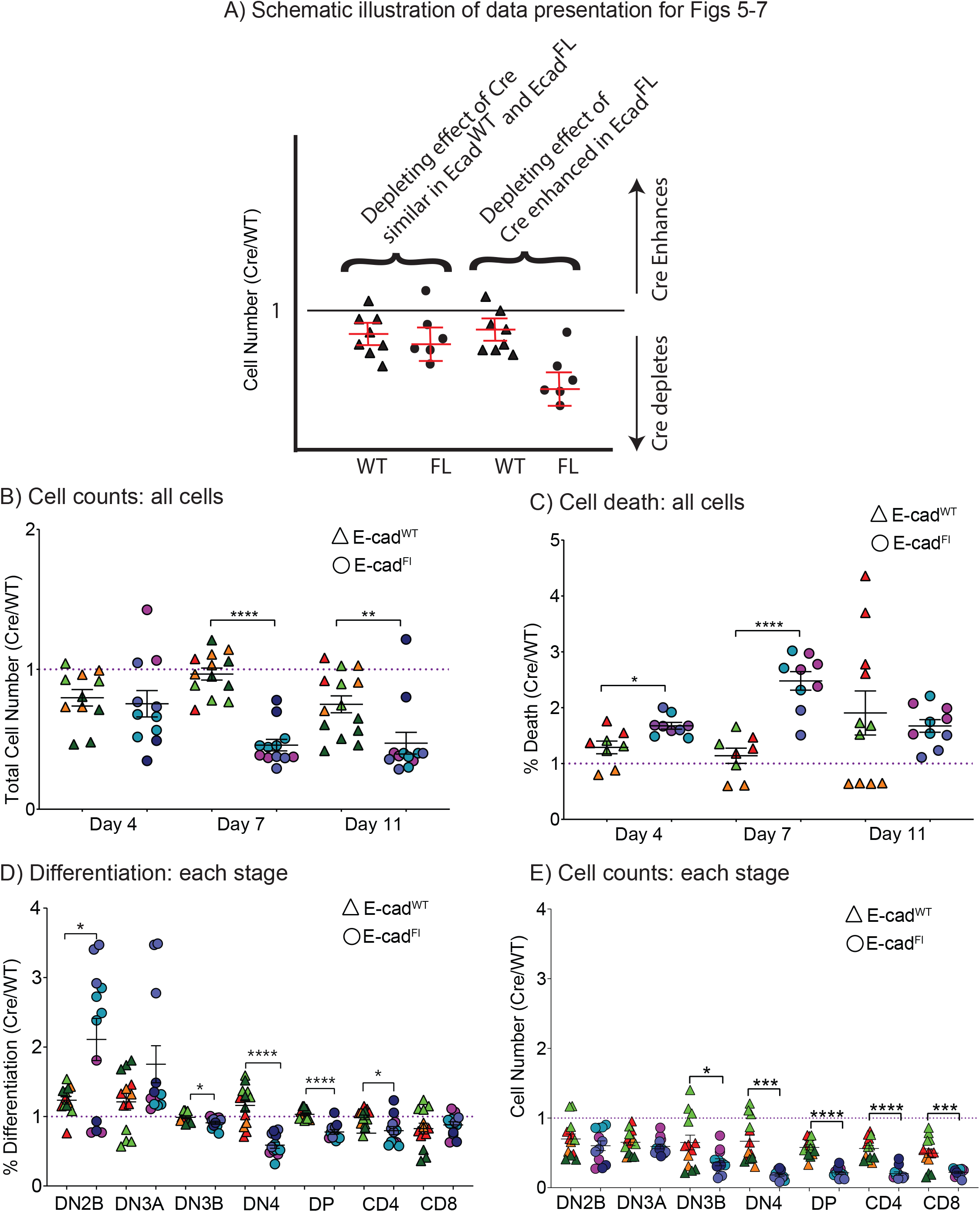
E-cadherin knockout effects T cell development. Developing T cells from E-cadherin^flox/flox^ (E-cad KO) or C57BL/6 (WT) mice were transfected with GFP or GFP-Cre on Day 4 and sorted on the basis of GPP expression on Day 8. GFP+ cells were re-seeded onto OP9-DL1 stromal cells and analysed for development on Day 4, 7 and 11. (A) Schematic of the organisation and interpretation of the data within the graphs. In a hypothetical scenario in which E-cadherin had no effect (LHS), both ratios would tend below the value of 1, due to Cre-mediated toxicity. In a hypothetical scenario in which E-cadherin had an effect, the ratios would shift from just below 1 downwards if E-cadherin further depleted cells, and upwards if E-cadherin enhanced cell numbers. E-cadherin knockout reduced (B) cell number and (C) at earlier time points it increased cell death. E-cadherin knockout reduced T cell development, as indicated by a reduction in (D) percent and (E) and number of cells that developed to DN3b onwards. Data is from 3 - 4 independent experiments, triangles indicate the E-cad^WT^ data and circles indicate E-cad^FL^ data; within the plots each dot represents a sample and the colour coding indicates the experiment. All data represented as Cre/WT, values less than 1 indicate Cre is greater, values above 1 indicate that WT was greater. * p < 0.05, ** p < 0.01, *** p< 0.001, ****p < 0.0001.

Inhibiting E-cadherin caused a large drop cell number and an increase in cell death at the earlier time points (**Fig. 5B, C**). E-cadherin knockout also hindered T cell differentiation, as evidenced in a reduction in the proportion of cells that progressed to DN3b and beyond, which was especially clear on Day 7 (**Fig. 5D, E and S8**). Intriguingly, the effect on cell number was most striking at the stages following β-selection despite the lack of expression at these later stages. These data indicate that E-cadherin is not required for survival during β-selection, but for the subsequent differentiation. Deleting E-cadherin also reduced the total and surface level of TCRβ (**Fig. S9**). Hence, E-cadherin is required for successful passage of DN3 cells through β-selection.

The loss of E-cadherin appeared to hinder T cell development specifically during β-selection, consistent with its expression at the DN3a and DN3b stage (Fig. 1g). To pinpoint more precisely the stage at which E-cadherin exerts its effects, we sorted for DN3a and DN3b cells and analysed their subsequent development up to 11 days after reseeding onto OP9-DL1 stromal cells. For starting populations of both DN3a and DN3b, cell number was greatly reduced in the absence of E-cadherin (**Fig. 6A, B**). The level of cell death and proliferation was not affected (**Fig. 6C-F)**, however, differentiation from both DN3a and DN3b to later stages was clearly hindered and this was especially evident on Day 7 (**Fig. 6G, H and S10**). These data, combined with the previous observation (Fig. 5) that E-cadherin deletion in unsorted cells impeded progression from DN3b onwards, indicates that both DN3a and DN3b cells need E-cadherin for optimal T cell development. Overall, we have shown that E-cadherin controls the regulation of division in developing DN3a T cells; the loss of E-cadherin caused misalignment of the mitotic spindle and its associated proteins, which delayed progression through the checkpoint. This affected the balance of cell fates after progression through the checkpoint which culminated in a reduction in the differentiation of the developing T cells.

**Figure 6.**
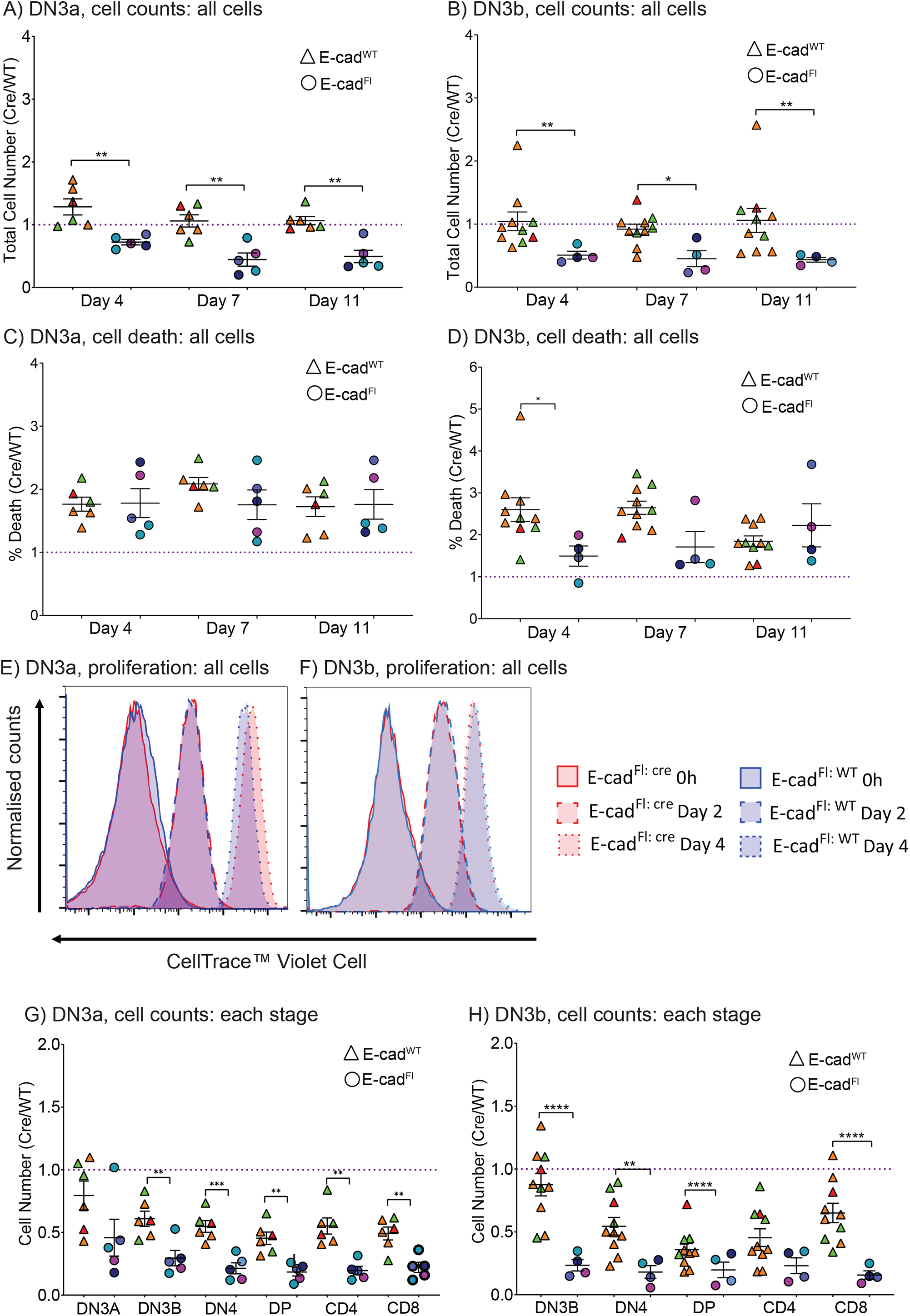
E-cadherin knockout hinders development of DN3a and DN3b T cells. Developing T cells from E-cadherin^flox/flox^ (E-cad KO) or C57BL/6 (WT) mice were transfected with GFP or GFP-Cre on Day 4 and sorted GPP+ DN3a or DN3b on Day 8. Cells were re-seeded onto OP9-DL1 stromal cells and analysed for cell death, proliferation and development. E-cadherin knockout reduced (A, B) cell number when the starting population was either DN3a or DN3b cells but did not affect cell death (C, D). (E, F) The cells were stained with Cell Trace and analysed after 2 and 4 days of culture. E-cadherin knockout had no effect on the rate of proliferation at any of the time points analysed. (G, H) E-cadherin knockout reduced T cell development, as indicated by a reduction total number of cells that developed to DN3b onwards. Data is from 3 -4 independent experiments, triangles indicate the E-cad^WT^ data and circles indicate E-cad^FL^ data; within the plots each dot represents a sample and the colour coding indicates the experiment. All data represented as Cre/WT, values less than 1 indicate Cre is greater, values above 1 indicate that WT was greater, * p < 0.05, ** p < 0.01, *** p< 0.001, ****p < 0.0001.

### Effect of E-cadherin signalling on T cell development was independent of Notch signalling

Notch is essential for progression through β-selection and subsequent development and also dictates the polarisation of Numb and α-adaptin in dividing DN3a cells (Ciofani and Zuniga-Pflucker 2005, Charnley, Ludford-Menting et al. 2020). We therefore hypothesised that the impact of E-cadherin deletion on spindle orientation might enhance the phenotype observed when Notch signalling is inhibited. To assess this, we combined E-cadherin deletion with pharmacological inhibition of Notch signalling. Inhibiting Notch slightly decreased the number of cells arising from a DN3a starting population as compared to E-cadherin-depleted cells that were not inhibited for Notch. This suggested an additional toxic effect of inhibiting Notch and raised the possibility of a functional interaction between E-cadherin and Notch restricted to the β-selection stage. However, this was not explained by changes in the number of apoptotic cells. The reduced total cell count was also not explained by a difference in cell count for any single differentiation stage between cells in which E-cadherin and Notch signalling had been inhibited versus cells where only E-cadherin was deleted (**Fig. 7**). These data indicate that the effects of E-cadherin on β-selection are not amplified by inhibition of Notch.

**Figure 7.**
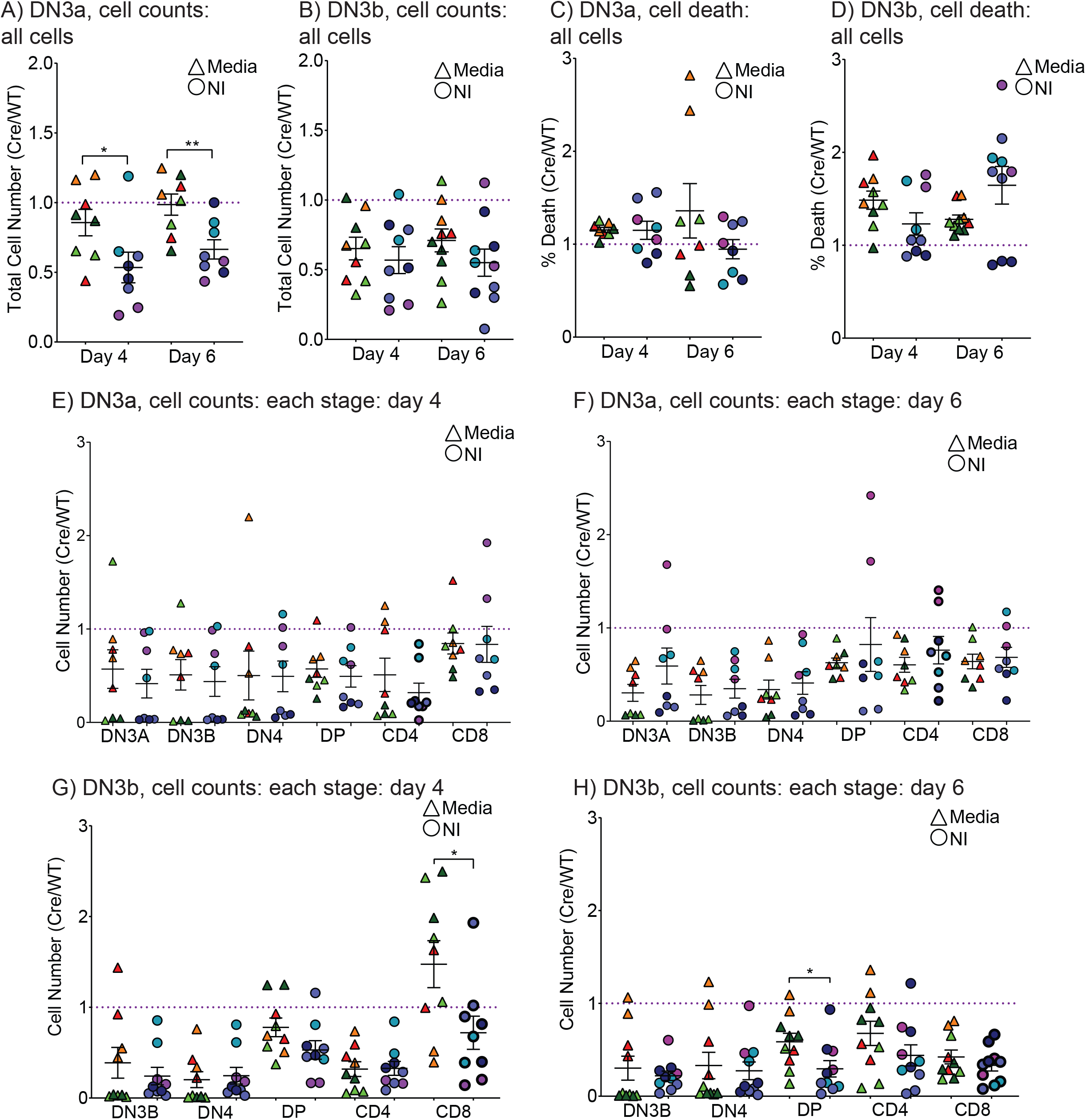
The effect of E-cadherin on developing T cell differentiation was independent of Notch signalling. Developing T cells from E-cadherin^flox/flox^ mice were transfected with GFP or GFP-Cre on Day 4 and sorted for GPP+ DN3a or DN3b cells on Day 8. Cells were re-seeded onto OP9-DL1 stromal cells, treated with either media or notch inhibitor and analysed for development on Day 4, 6. The combination of deleting E-cadherin and inhibiting Notch signalling (A) slightly reduced cell number when DN3a cells were used as the starting population. Inhibiting notch signalling didn’t effect (B) cell death or development, regardless of whether the starting population was (C) DN3a or (D) DN3b. Data from is 3 - 4 independent experiments, triangles indicate samples treated with media and circles indicate samples treated with the notch inhibitor; within the plots each dot represents a sample and the colour coding indicates the experiment, All data represented as Cre/WT, * p < 0.05, ** p < 0.01, *** p< 0.001.

**Figure 8:**
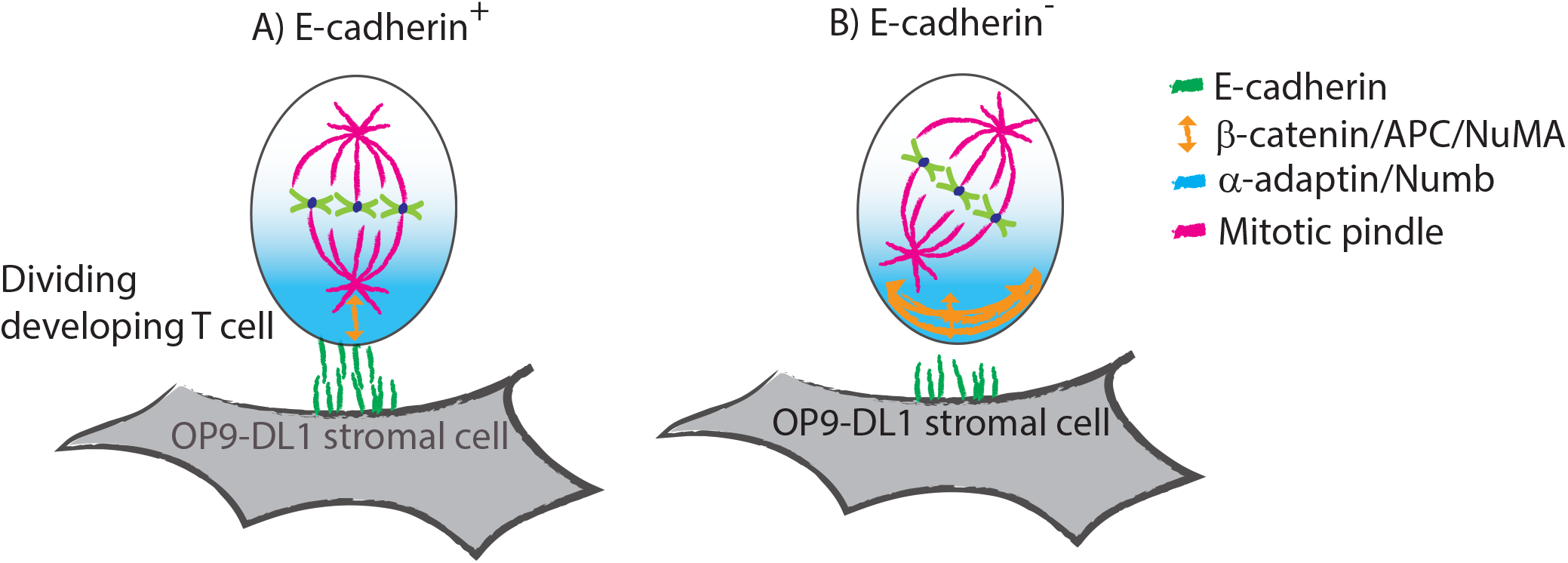
A model of the role of E-cadherin in controlling spindle orientation in dividing T cells. (A) E-cadherin ligation of E-cadherin presented by the OP9-DL1 stromal cell, or E-cadherin functionalised surfaces, enables spindle orientation proteins, including β-catenin, APC and NuMA, to align the mitotic spindle with the axis of polarity of cell fate proteins α-adaptin and Numb. (B) In the absence of endogenous E-cadherin in the developing T cell, alignment of the mitotic spindle is disrupted. This is associated with delayed progression through the β-selection checkpoint and subsequently reduced T cell differentiation.

## Discussion

It is now clear that TCR signalling during β-selection shapes the TCR repertoire (Mallis, Bai et al. 2015, Bovolenta, García-Cuesta et al. 2022, Duke-Cohan, Akitsu et al. 2022). Unlike in T cells, TCR signalling during β-selection is hampered by the lack of the TCRα chain and coreceptors, CD4 and CD8. A possible means of facilitating this handicapped TCR signal is the provision of extra adhesion molecules in the form of a niche (Ciofani, Schmitt et al. 2004, Mallis, Bai et al. 2015, Zhao, Yoganathan et al. 2019, Allam, Charnley et al. 2021). In particular, Notch and CXCR4 can facilitate this process via synapse formation (Allam, Charnley et al. 2021) and by coordinating polarity during division (Pham, Shimoni et al. 2015, Charnley, Ludford-Menting et al. 2020). However, we lack a comprehensive understanding of the range of cues that can support the low affinity interaction between pre-TCR and MHC to drive developing T cells through β-selection. E-cadherin is well-known mediator of cell–cell junctions which controls multiple aspects of cell behaviour, including adhesion and synapse formation (Perez-Moreno, Jamora et al. 2003, Bamji 2005, Halbleib and Nelson 2006, Martin-Belmonte and Perez-Moreno 2011, Martín-Cófreces, Vicente-Manzanares et al. 2018), polarity (Charnley, Kroschewski et al. 2012) and division (den Elzen, Buttery et al. 2009). E-cadherin influences downstream fate decisions in many aspects of development and has long been known as a central player in communication from tissue organisation to cell signalling (Jamora and Fuchs 2002, Yap, Duszyc et al. 2018). This coupled with the upregulation of E-cadherin specifically at this stage of T cell development (Lee, Sharrow et al. 1994, Muller, Luedecker et al. 1997) motivated us to examine the role of E-cadherin in T cell development.

To mimic the presentation of E-cadherin in the niche and isolate its effect on developing T cells we used functionalised surfaces coated with E-cadherin (Charnley, Kroschewski et al. 2012). The exogenous presentation of E-cadherin supported cell attachment and polarisation of cell fate determinants to the cell surface interface. Endogenous E-cadherin within the developing T cell was recruited to the cell contact and increased the stability of the immunological synapse. Thus, E-cadherin formed homotypic E-cadherin-E-cadherin interactions which promoted the formation and stability of the synapse. We have previously shown that Notch and CXCR4 coordinated to form a synapse between the developing T cells and OP9-DL1 stromal cells, which facilitates progression through β-selection (Charnley, Ludford-Menting et al. 2020, Allam, Charnley et al. 2021). We have now expanded this to show that E-cadherin also facilitates this interaction. Given the known role of E-cadherin in cell-cell interactions, this might suggest a role for E-cadherin in reinforcing the weak pre-TCR-MHC interactions to support TCR signalling and progression through β-selection.

The role of E-cadherin was not restricted to supporting the adhesion of the developing T cell. In dividing T cells endogenous E-cadherin was polarised and regulated the distribution of three proteins, β-catenin, adenomatous polyposis coli (APC) and NuMA, that control the spindle orientation. How might E-cadherin affect the distribution of these regulators of the mitotic spindle? In various organisms, the position of the mitotic spindle is controlled by cortical force generator complexes, such as the LGN/NuMA/Gα complex, interacting with dynein and astral microtubules (Matsuzaki and Shitamukai 2015, di Pietro, Echard et al. 2016). In epithelial cells E-cadherin acts as an instructive cue to direct the assembly of NuMA/LGN complex at the cell-cell contact (Gloerich, Bianchini et al. 2017). This couples the cell division axis with the intracellular adhesion by stabilising the interaction between cortical cues at the cell contact and the astral microtubules. β-catenin directly binds to E-cadherin and also interacts indirectly with actin and microtubule networks (Shahbazi and Perez-Moreno 2015), while APC binds to β-catenin and microtubule binding proteins (Yamashita, Jones et al. 2003, Yamashita 2010). Hence, all of these proteins act as a link between E-cadherin and the mitotic spindle. These findings suggest that disrupting E-cadherin prevented it from acting as an instructive cue, which altered the localisation of β-catenin, NuMa and APC and hindered the correct spindle alignment. Interestingly, loss of APC has previously been shown to disrupt thymic development and slow proliferation due to a delayed exit from mitosis (Gounari, Chang et al. 2005). This was proposed to be due to activation of the mitotic checkpoint, which delays entry into anaphase until there is proper alignment of the chromosomes at the spindle (Gounari, Chang et al. 2005, Charnley, Anderegg et al. 2013). In this system we observed a similar increase in the time require to complete division, indicating that spindle misorientation delayed division. Thus, E-cadherin is required for the proper recruitment of these proteins to the cell contact, which facilities the correct alignment of the mitotic spindle and progression through division.

E-cadherin has been shown to link external cues with mitotic spindle orientation and asymmetric cell division (ACD) in many developmental systems (Le Borgne, Bellaïc he et al. 2002, den Elzen, Buttery et al. 2009, Neumüller and Knoblich 2009, Martin-Belmonte and Perez-Moreno 2011, Werts and Goldstein 2011, Gloerich, Bianchini et al. 2017). Intriguingly, despite the capacity of exogenous E-cadherin to promote polarity in dividing T cells, endogenous E-cadherin was not required for polarisation *per se* of specific cells fate determinants, namely Numb and α-adaptin (Fig 2). This suggests a redundancy in the cues that drive polarisation. This redundancy is in contrast to Notch and CXCR4, which cooperate to drive the polarisation of these cell fate determinants, and VCAM-1, which reduced polarisation (Charnley, Ludford-Menting et al. 2020). Thus, it appears that Notch and CXCR4 signalling are the critical drivers for the polarisation of cell fate determinants and E-cadherin plays a more important role in coordinating the orientation of the mitotic spindle with the axis of polarity. Thus, multiple external cues can coordinate to dictate how the developing T cell divides and develops.

Another non-mutually exclusive mechanism by which E-cadherin may mediate the disruption to β-selection might relate to the well-known role of Wnt signalling in T cell development (Gounari and Khazaie 2022). A central effector of the canonical Wnt pathway is β-catenin, which is stabilized by extracellular Wnt ligands to enable binding to TCF/LEF in the nucleus (Mosimann, Hausmann et al. 2009). β-catenin is not essential for T cell development, but plays important roles during β-selection in avoiding oncogenic transformation, with stabilization of β-catenin leading to several T cell malignancies (Gounari and Khazaie 2022). Importantly for this study, another critical layer of control of β-catenin is E-cadherin, with cadherin-mediated adhesion regulating β-catenin localisation and signalling (Heuberger and Birchmeier 2010). Thus, another possible explanation for the effects of E-cadherin deletion is that E-cadherin-mediated adhesion, through β-catenin, might regulate TCF/LEF signalling during β-selection.

In an interesting twist, the Wnt pathway is also considered a key regulator of planar cell polarity (PCP) and its control of asymmetric cell division (Segalen and Bellaiche 2009). Although PCP is often β-catenin-independent (Phillips and Kimble 2009) in the nematode *Caenorhabditis elegans*, asymmetric activity of Wnt signalling is dictated by asymmetric localisation of β-catenin at cell division (Sugioka, Mizumoto et al. 2011). Conversely, immobilised Wnt signals have been demonstrated to drive asymmetric distribution of β-catenin and APC during division in embryonic and human skeletal stem cells (Habib, Chen et al. 2013, Lowndes, Rotherham et al. 2016, Okuchi, Reeves et al. 2021). Our finding here that β-catenin was asymmetric during DN3a cell division suggests the possibility that E-cadherin might control Wnt signalling in part by dictating asymmetric recruitment of β-catenin to one daughter during β-selection. Further exploration is needed to determine whether polarity of β-catenin, APC and NuMA during division plays an important role in β-selection, and how E-cadherin influences this pathway.

To determine whether there was correlation between the role of E-cadherin in supporting adhesion and division during β-selection and the subsequent differentiation in T cells we examined whether E-cadherin was required for T cell development. T cell development was indeed hindered from DN3b onwards, indicating that E-cadherin is required for T cell differentiation. This developmental defect in E-cadherin deficient T cells might be explained by one or more these mechanisms; namely an alteration in binding to the β-selection niche; a defect on the alignment of the mitotic spindle with the polarity axis during division; and/or a defect in Wnt signalling. Overall, this works provides new insights into the role of E-cadherin in the β-selection stage of T cell development.

## Methods

### Primary developing T cell co-culture

Mouse hematopoietic stem cells (isolated from C57BL/6 or E-cadherin flox/flox fetal livers at E14.5) were seeded onto OP9-DL1 stromal cells (received from Juan-Carlos Zúñiga-Pflücker, University of Toronto, Sunnybrook Health Sciences Centre, Toronto, Ontario, Canada) at a 1:1 ratio in 6 well plates (2 × 10^5^) in Minimal Essential Medium Alpha Modification supplemented with foetal calf serum (10% v/v), glutamine (1 mM), β-mercaptoethanol (50 μM), sodium pyruvate (1 nM), HEPES (10 mM), penicillin/streptomycin (100 ng/mL), mouse interleukin 7 (IL-7, 1 ng/mL) and mouse FMS-like tyrosine kinase 3 (Flt-3, 5 ng/mL). Hematopoietic cells were harvested via forceful pipetting and co-cultured on fresh OP9-DL1 stromal cells every 3–4 days. All mice were maintained in a specific pathogen-free environment with food and water freely available. All experiments on mice were performed in accordance with the Animal Experimentation Ethics Committee of La Trobe University (Ethics approval number AEC20015).

### Retroviral transduction

Phoenix E cells (provided by Garry Nolan, Stanford University, Stanford, CA) were maintained at 37°C and at 10% CO_2_ in Dulbecco’s Minimal Essential Medium supplemented with foetal calf serum (10% v/v), L-glutamine (1 mM) and penicillin/streptomycin (100 ng/mL). Calcium phosphate transfection was performed on Phoenix E cells with 15-20 μg of the following pMSCV retroviral constructs: GFP, GFP-Cre, Cherry-Numb and Cherry-α-adaptin in 10 cm dishes (Corning). Viral supernatant was harvested 48 h after transfection and added to 6 well plates that had been precoated with 15 μg/ml RetroNectin (Takara Bio Inc.) and blocked with 2% BSA. After addition of the viral supernatant, plates were spun at 2,000 *g* for 1 h and incubated for 1 h at 37°C. 5 × 10^5^ hematopoietic cells (day 4 co-culture) were added, and plates were spun for 1 h at 1,200 *g*.

### Flow cytometry

Developing T cell subsets were purified by staining for the cell surface markers CD44 and CD25 to discriminate between DN 1-4, CD28 to discriminate between DN3a and DN3b, and CD4 and CD8 to discriminate between the DN and DP populations. Hematopoietic cells were harvested from OP9-DL1 cocultures by forceful pipetting and stained on ice for one hour with the following antibodies: Pacific Blue CD4 (1:500; BD Pharmingen, 558107), BV711 CD8a (1:1000; BD Biosciences, 563676), BV510 CD44 (1:500; BD Biosciences, 563114), BV786 CD25 (1:500; BD Biosciences, 564023), Pe/Cy7 CD28 (1:200; Biolegend, 122014) for one hour on ice prior to sorting. DN3a cells were isolated on a FACS (FACS Aria III; BD Biosciences) based on surface expression (CD28^lo^/CD25^+^/CD44^lo^/CD4^−^/CD8^−^). Retrovirally transduced cells were sorted 72 hours after transduction and also sorted on the basis of GFP and Cherry fluorescence. For the analysis of T cell differentiation, cells were additionally stained with PE E-cadherin (1:400, Biolegend, 147304) or Alexa Fluor® 647 E-cadherin (1:300, Biolegend, 147308) and 7-AAD viability stain (0.25 μg; BD Biosciences, 559925).

For the analysis of surface expression of TCRβ the cells were stained with PE CD4 (1:600; eBioscience, 12-0041-81), APC-eFluor 780 CD8a (1:300, eBioscience, 47-0081-82), PerCP/Cy5.5 CD44 (1:300, Biolegend 103032), BV510 CD25 (1:500, BD Biosciences, 563037), Pe/Cy7 CD28 (1:200, Biolegend, 122014) and APC TCRβ (1:400, Biolegend, 109212). To analyse the surface expression of αE and β7 the cells were stained with BV605 CD4 (:500; BD Pharmingen, 563151), BV711 CD8, BV510 CD44, BV786 CD25, PeCy7 CD28, APC E-cad, PE αE (eBioscience, 12-1031-82, 1:200), biotin β7 (eBioscience, 13-5867-82, 1:500). For biotin conjugated antibodies (β7) the DN3a cells were rinsed and stained with BV421 Streptavidin (1:600, BD Biosciences, 563259) for one hour on ice prior to analysis.

### Intracellular staining

Total protein expression level of TCRβ was determined using Intracellular Fixation and Permeabilization Buffer Set (eBioscience, 88-8824). Briefly, developing T cells were harvested and stained with surface markers for CD4/CD8/CD44/CD25/CD28 as per setup used for surface expression of TCRβ, then fixed for 30 mins at room temperature in fixative buffer, then washed twice in distilled water supplemented with 10% permeabilization buffer. Fixed developing T cells were resuspended with APC TCRβ (1:400, Biolegend, 109212) in 10% permeabilization buffer at room temperature for 1 hr, washed twice and resuspended in culturing media for flow cytometry analysis.

### Substrate functionalisation

For live cell imaging PDMS cell paddocks (120 μm x 120 μm) (Day, Pham et al. 2009) were rendered hydrophilic by exposure to air plasma at 1.5 × 10^-2^ mbar for 10 minutes and placed into an 8 well chamber slide (Ibidi) prior to protein functionalisation or OP9-DL1 stromal cell culture. PDMS cell paddocks or 24 well plates were coated with PA (235 nM, 1 hour), rinsed with PBS and further functionalised, if required, with Fc-E-cadherin (72 nM; R&D Systems, RDS748EC050, PBS plus 0.2 mM CaCl_2_) or Fc-DL4 (86 nM; BioVision, P1163), for 30 minutes. The substrates were then washed in PBS (including 2 mM CaCl_2_ for Fc-E-cadherin coatings) and media before cell deposition.

### Proliferation and development

For the analysis of the effect of disrupting E-cadherin on differentiation, 24 well plates were cultured OP9-DL1 stromal cells. 1 × 10^5^ GFP^+^ or GFP-Cre^+^ or DN3a (GFP^+^ or GFP-Cre^+^) versus DN3b (GFP^+^ or GFP-Cre^+^) cells were seeded into the wells and assessed at 4, 7 and 11 days by flow cytometry. Proliferation was assessed by incubating the cells with CellTrace™ Violet Cell (C34571, Invitrogen, 5 μM) for 20 minutes prior to seeding the cells onto OP9-DL1 stromal cells. Proliferation was assessed on Day 2 and 4 using flow cytometry. To disrupt Notch signalling during T cell development we used the γ-secretase inhibitor, dibenzazepine (DBZ; 0.05 μM) (Charnley, Ludford-Menting et al. 2020). Purified DN3a (GFP^+^ or GFP-Cre^+^) versus DN3b (GFP^+^ or GFP-Cre^+^) cells were co-cultured on OP9-DL1 stromal cells in the presence or absence of DBZ and assessed by flow cytometry.

### Immunofluorescence and fixed image acquisition by confocal microscopy

2 × 10^4^ DN3a cells were added to 8 well chamber slide (Thermo Fisher Scientific) precoated with PA plus Fc-E-cadherin or OP9-DL1 stromal cells and cultured for 2 or 15 hours. For stromal cell-free culture cells were cultured in the presence of Flt-3 (5 ng/mL), CXCL12 (10 nM) and IL-7 (10 ng/ml) (Janas, Varano et al. 2010, Charnley, Ludford-Menting et al. 2020). Cells were then fixed, permeabilized, and labelled with primary antibodies for α-tubulin (1:800; Rocklands, 600-401-880 or 200-301-880) and test proteins, α-adaptin (1:200; Abcam, Ab2730), Numb (1:200; Abcam, Ab4147), CD25 (1:1000; BD Pharmingen, 553070), β-catenin (1:500, Sigma-Aldrich, C2206), NuMA (1:1000, Abcam, ab109262), APC (1:200, gift from I. Nathke, University of Dundee) and the appropriate secondary antibodies and then mounted with Prolong gold antifade with DAPI (Molecular Probes, P36930). The slides were examined using an Olympus FV3000 microscope (Olympus) and 60x objective lens (1.35 NA Oil Plan Apochromat). 3D images of the cells were acquired with a z distance of 0.5 μm and maximum intensity projections of z sections created using ImageJ. For quantifications, the investigator was blind to the sample allocation at the moment of scoring. For the analysis of polarisation during interphase α-tubulin staining was used to identify cells with the MTOC at the cell-cell / cell-substrate interface and the cell was scored depending on whether the protein of interest was diffuse or enriched at the MTOC.

To determine polarisation during division mitotic cells were identified by the presence of a mitotic spindle and the cell division was assigned as symmetric if the fluorescence of the test protein was evenly distributed between the two daughter cells and asymmetric if the fluorescence was greater in one of the daughter cells than the other daughter cell. To determine if inhibiting E-cadherin caused a change in the spatial distribution of β-catenin, APC and NuMA in developing T cells, the fluorescent intensity at the cortex, midbody and centromeres, which were identified as the key regions that these proteins polarised to during division (Olmeda, Castel et al. 2003, den Elzen, Buttery et al. 2009, Poulson and Lechler 2010, Gloerich, Bianchini et al. 2017, Kiyomitsu and Boerner 2021), were measured and ratioed to the fluorescent intensity of the cytoplasm. A high relative concentration indicates that the protein is concentrated in the cortex, midbody and centromeres, while a low relative concentration indicates that the relative level of the protein in the cytoplasm is increased.

### Live cell image acquisition and analysis

2 × 10^4^ DN3a cells were added to 8 well chamber slide (Ibidi) with the PDMS cell paddocks precoated with PA plus Fc-linked proteins or OP9-DL1 stromal cells. For stromal cell-free culture cells were cultured in the presence of Flt-3 (5 ng/mL), CXCL12 (10 nM) and IL-7 (10 ng/mL) (Janas, Varano et al. 2010, Charnley, Ludford-Menting et al. 2020). Images were captured on a spinning disc confocal system fitted with an inverted microscope and temperature-controlled chamber maintained at 37°C and 5% CO_2_. Images are acquired using a 40x air objective (0.95 NA) and multiple stage positions are recorded every 3 min for 20 hours, with 8-slice z-stacks of 1 μm thickness. As described in (Oliaro, Van Ham et al. 2010, Shimoni, Pham et al. 2014, Pham, Shimoni et al. 2015, Charnley and Russell 2017, Charnley, Ludford-Menting et al. 2020), the polarisation of fluorescent markers in the nascent daughter cells was quantified by measuring the total fluorescence intensity of each daughter cell and applying a “Polarization Ratio” (PR) equation: PR = (ΣH1 − ΣH2)/ (ΣH1 + ΣH2); where the difference in total intensity between Daughter 1 (ΣH1) and Daughter 2 (ΣH2) is divided by the sum of intensities in both Daughter 1 (H1) and Daughter 2 (H2). A low polarisation ratio indicates that the protein was evenly distributed between the two nascent daughter cells, while a high polarisation ratio indicates that the protein was concentrated in one of the daughter cells. A PR of 0.17 was used as a cutoff to designate cells as undergoing ACD (PR > 0.17) or SCD (PR < 0.17). This cutoff was determined from the spread of the diffuse protein; specifically, this cut-off resulted in minimal number of the diffuse protein divisions being assigned as asymmetric. On the functionalised surfaces, only cells that had not interacted with another cell for at least 30 minutes prior to division were selected for analysis. This enabled the isolation of the effect of the immobilised protein on ACD. To assess the role of T cell – T cell interactions, developing T cells were cultured on PA coated surfaces and dividing cells that formed and maintained cell contacts for at least 30 minutes prior to and throughout division were selected for analysis. To compare the distribution of PR values between the different cell culture platforms, outliers (which were identified by their high PR diffuse values and consisted of less than 10% of dividing cells) were removed (Charnley and Russell 2017, Charnley, Ludford-Menting et al. 2020) and the data was re-plotted as violin plots using the kernel density function.

To assess the role of the orientation of the mitotic spindle relative to the polarity cue, the division angle was determined by measuring the angle between the interface between the DN3a cell and OP9-DL1 stromal cell and the long axis of the dividing cell. The division angle was plotted *versus* PR to visualise the correlation between the orientation of the mitotic spindle and level of polarisation. The stability of the immunological synapse was analysed in cells transfected with cherry-tubulin and either GFP or GFP-Cre. Cells were triaged for cells in which MTOC was recruited to the interface with an OP9-DL1 cell for at least 30 minutes and tracked the cells to determine the duration of the synapse.

## Statistical Analysis

All data was collected from 2 to 8 independent experiments. For scatter and bar plots data is shown as mean ± SEM, median value is indicated on the violin plots. The following tests were used for the statistical analysis: Student’s unpaired two-way t-test with unequal variance was used to determine the difference between mean values for the scatter plots and calculated using Microsoft Office Excel. The violin plots were analysed for differences in the distribution of the data using Two-sample Kolmogorov-Smirnov (KS) test (Young 1977) in Graphpad Prism software. Level of statistical significance is indicated as: *p < 0.05, **p < 0.01, ***p < 0.001.

## Acknowledgments

This work was done on Wurundjeri land of the Kulin nation, and we pay our respects to the Elders past and present. We thank Juan Carlos Zúñiga-Pflücker (University of Toronto, Sunnybrook Health Sciences Centre, Toronto, Canada), for OP9 stromal cell lines. This work was performed in part at the Biointerface Engineering Hub @ Swinburne, part of the Victorian node of the Australian National Fabrication Facility (ANFF), a company established under the National Collaborative Research Infrastructure Strategy to provide nano-and micro-fabrication facilities for Australia’s researchers.

