## Supplementary data for "E-cadherin in developing murine T cells controls spindle alignment and differentiation during β-selection"

**Supplementary figure legends**

**Figure S1. DN3a cells undergo polarised divisions on E-cadherin functionalised surfaces**

To determine whether exogenous E-cadherin could trigger polarisation and asymmetric cell division DN3a cells were seeded onto PA + E-cadherin coated surfaces and either fixed and stained after 15 hours or imaged for 20 hours using time lapse microscopy. (A) DN3a cells attached to the E-cadherin surfaces and was comparable to level of attachment previously observed on protein A (PA) only surfaces (Charnley, Ludford-Menting et al. 2020). Representative images (left) and PR scatter plots (right) for dividing DN3a cells transduced with α-adaptin and GFP (diffuse control) and cultured on (B) OP9-DL1 stromal cells (C) PA with cell-cell contact (D) PA + E-cadherin or (E) PA alone. PR was calculated as the difference in fluorescence between the two halves divided by the sum of fluorescence. The blue shaded region indicates divisions that were assigned as asymmetric when the 0.17 cut-off was applied. The distribution of PR values was greater in the presence of E-cadherin. (F) The expression of αEβ7, which is an alternative binding partner for E-cadherin, was assessed in developing T cells using flow cytometry. Developing T cells express β7 at the DN2b stage of development and expression decreased at later stages of development. A low level of expression for αE was observed at all stages of development. Flow cytometry plots of one representative experiment are shown, n = 4 independent experiments, scale bar = 10 µm.

**Figure S2. Endogenous E-cadherin was required for polarisation of α-adaptin in dividing DN3a cells cultured on E-cadherin functionalised surfaces**

Developing T cells from E-cadherin$^{flox/flox}$ mice were transfected with GFP or GFP-Cre on Day 4 and sorted on the basis of GPP expression on Day 8. GFP+ cells were re-seeded onto OP9-DL1 stromal cells and analysed for development on Day 4, 7 and 11. (A) E-cadherin knockout was confirmed in DN3 cells after sorting and this was maintained over the course of the experiment. To assess the role of endogenous E-cadherin in polarisation during division DN3a cells were transduced with GFP (E-cad$^{+}$) or GFP-Cre (E-cad$^{-}$) and α-adaptin were seeded onto functionalised surfaces and imaged for 20 hours using time lapse microscopy. Representative images (left) and PR scatter plots (right) for dividing DN3a cells (B) E-cad$^{+}$ or (C) E-cad$^{-}$ on DL4 surfaces, (D) E-cad$^{+}$ or (E) E-cad$^{-}$ on E-cadherin coated surfaces or (F) E-cad$^{-}$ on PA. E-cadherin knockout reduced the level of ACD on E-cadherin coated surfaces. n = 6-7 independent experiments.

**Figure S3. Endogenous E-cadherin was not required for the polarised distribution of cell fate proteins**

(A) OP9-DL1 expression of E-cadherin expression was confirmed using flow cytometry. Flow cytometry plots of one representative experiment are shown, n = 3 independent experiments. (B – E) Developing T cells from E-cadherin$^{flox/flox}$ mice were transfected with GFP (E-cad$^{+}$) or GFP-Cre (E-cad$^{-}$) and cherry-α-adaptin or cherry-Numb on Day 4 and sorted on Day 8. DN3a cells were re-seeded onto OP9-DL1 stromal cells and imaged for 20 hours using time lapse microscopy. Representative images (left) and PR scatter plots (right) for dividing DN3a cells (B) E-cad$^{+}$ and α-adaptin, (C) E-cad$^{-}$ and α-adaptin, (D) E-cad$^{+}$ and Numb and (E) E-cad$^{-}$ and Numb and seeded onto OP9-DL1 cells. n = 6-7 independent experiments

**Figure S4. Distribution of β-catenin in dividing DN3a cells and the role of endogenous E-cadherin**

To determine the role of endogenous E-cadherin in the distribution of β-catenin in dividing DN3a cells developing T cells from E-cadherin$^{flox/flox}$ mice were transfected with GFP (E-cad$^{+}$) or GFP-Cre (E-cad$^{-}$). GFP+ cells were re-seeded onto OP9-DL1 stromal cells and fixed and stained after 15 hours. Representative images of the distribution of β-catenin during division in A) E-cad$^{+}$ DN3a cells and B) E-cad$^{-}$ cells, bar = 10 µm.

**Figure S5. Distribution of adenomatous polyposis coli (APC) in dividing DN3a cells and the role of endogenous E-cadherin**

To determine the role of endogenous E-cadherin in the distribution of adenomatous polyposis coli (APC) in dividing DN3a cells developing T cells from E-cadherin$^{flox/flox}$ mice were transfected with GFP (E-cad$^{+}$) or GFP-Cre (E-cad$^{-}$). GFP+ cells were re-seeded onto OP9-DL1 stromal cells and fixed and stained after 15 hours. Representative images of the distribution of adenomatous polyposis coli (APC) during division in A) E-cad$^{+}$ DN3a cells and B) E-cad$^{-}$ cells, bar = 10 µm.

**Figure S6. Distribution of NuMA in dividing DN3a cells and the role of endogenous E-cadherin**

To determine the role of endogenous E-cadherin in the distribution of NuMA in dividing DN3a cells developing T cells from E-cadherin$^{flox/flox}$ mice were transfected with GFP (E-cad$^{+}$) or GFP-Cre (E-cad$^{-}$). GFP+ cells were re-seeded onto OP9-DL1 stromal cells and fixed and stained after 15 hours.

Representative images of the distribution of NuMA during division in A) E-cad$^{+}$ DN3a cells and B) E-cad$^{-}$ cells, bar = 10 µm.

**Figure S7. E-cadherin knockout effects T cell development**

Developing T cells from E-cadherin$^{flox/flox}$ mice were transfected with GFP (E-cad$^{+}$) or GFP-Cre (E-cad$^{-}$) on Day 4 and sorted on the basis of GPP expression on Day 8. GFP+ cells were re-seeded onto OP9-DL1 stromal cells and analysed for development on Day 4, 7 and 11. Deleting E-cadherin reduced (A) cell number and (B) increased cell death. E-cadherin knockout reduced T cell development, as indicated by a reduction in (C) percent and (D) and total number of cells that developed to DN3b onwards. All data represented as mean ± SEM, * p < 0.05, ** p < 0.01, *** p< 0.001.

**Figure S8. E-cadherin knockout effects T cell development**

Developing T cells from E-cadherin$^{flox/flox}$ (E-cad KO) or C57BL/6 (WT) mice were transfected with GFP (E-cad$^{+}$) or GFP-Cre (E-cad$^{-}$) on Day 4 and sorted on the basis of GPP expression on Day 8. GFP+ cells were re-seeded onto OP9-DL1 stromal cells and analysed for development on Day 4, 7 and 11. E-cadherin knockout reduced T cell development, as indicated by a reduction in (A) percent and (B) and number of cells that developed to DN3b onwards. Data is from 3 - 4 independent experiments, triangles indicate the E-cad$^{WT}$ data and circles indicate E-cad$^{FL}$ data; within the plots each dot represents a sample and the colour coding indicates the experiment. All data represented as Cre/WT, values less than 1 indicate Cre is greater, values above 1 indicate that WT was greater. * p < 0.05, ** p < 0.01, *** p< 0.001, ****p < 0.0001.

**Figure S9. E-cadherin knockout reduces TCRβ expression**

Developing T cells from E-cadherin$^{flox/flox}$ mice were transfected with GFP (E-cad$^{Fl:WT}$) or GFP-Cre (E-cad$^{Fl:Cre}$) on Day 4 and analysed on Day 10 to 12. E-cadherin knockout slightly reduced (A) total and (B) surface expression of TCRβ from DN3b onwards. Flow cytometry plots of one representative experiment are shown, n = 3 independent experiments.

**Figure S10. E-cadherin knockout effects development of DN3a and DN3b T cells**

Developing T cells from E-cadherin$^{flox/flox}$ (E-cad KO) or C57BL/6 (WT) mice were transfected with GFP (E-cad$^{+}$) or GFP-Cre (E-cad$^{-}$) on Day 4 and sorted for GPP+ (A) DN3a or (B) DN3b on Day 8. Cells were re-seeded onto OP9-DL1 stromal cells and analysed for cell death, proliferation and development. E-cadherin knockout reduced T cell development, as indicated by a reduction total number of cells that developed to DN3b onwards. Data is from 3 - 4 independent experiments, triangles indicate the E-cad$^{WT}$ data and circles indicate E-cad$^{FL}$ data; within the plots each dot represents a sample and the colour coding indicates the experiment. All data represented as WT/Cre, * $p < 0.05$, ** $p < 0.01$, *** $p< 0.001$, ****$p < 0.0001$.

Fig. S1

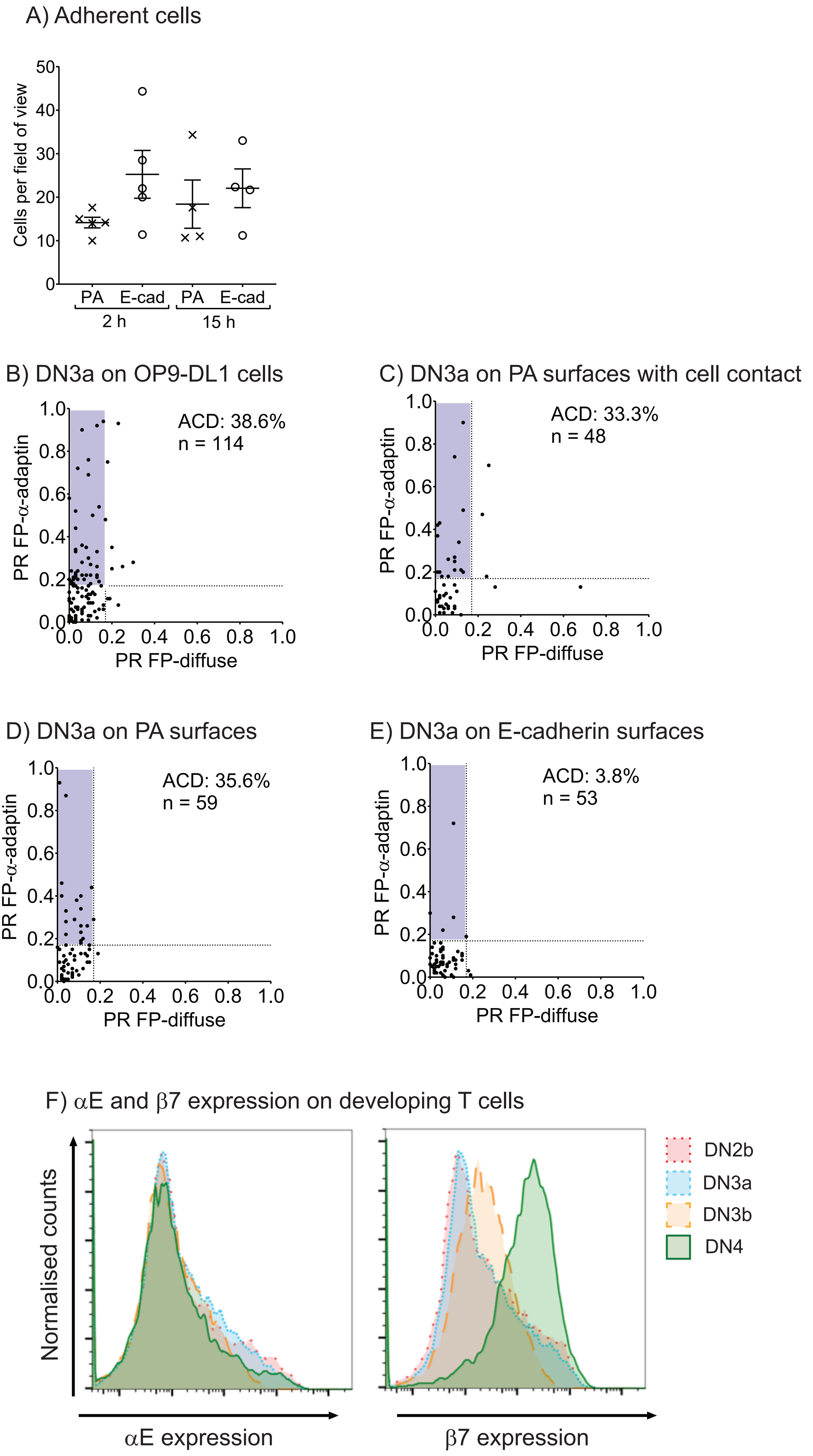

A) Adherent cells
Cells per field of view
50
40
30
20
10
0
PA
E-cad
PA
E-cad
2 h
15 h
B) DN3a on OP9-DL1 cells
ACD: 38.6%
n = 114
PR FP-α-adaptin
PR FP-diffuse
C) DN3a on PA surfaces with cell contact
ACD: 33.3%
n = 48
PR FP-α-adaptin
PR FP-diffuse
D) DN3a on PA surfaces
ACD: 35.6%
n = 59
PR FP-α-adaptin
PR FP-diffuse
E) DN3a on E-cadherin surfaces
ACD: 3.8%
n = 53
PR FP-α-adaptin
PR FP-diffuse
0.0
0.2
0.4
0.6
0.8
1.0
F) αE and β7 expression on developing T cells
Normalised counts
αE expression
β7 expression
DN2b
DN3a
DN3b
DN4

Fig. S2

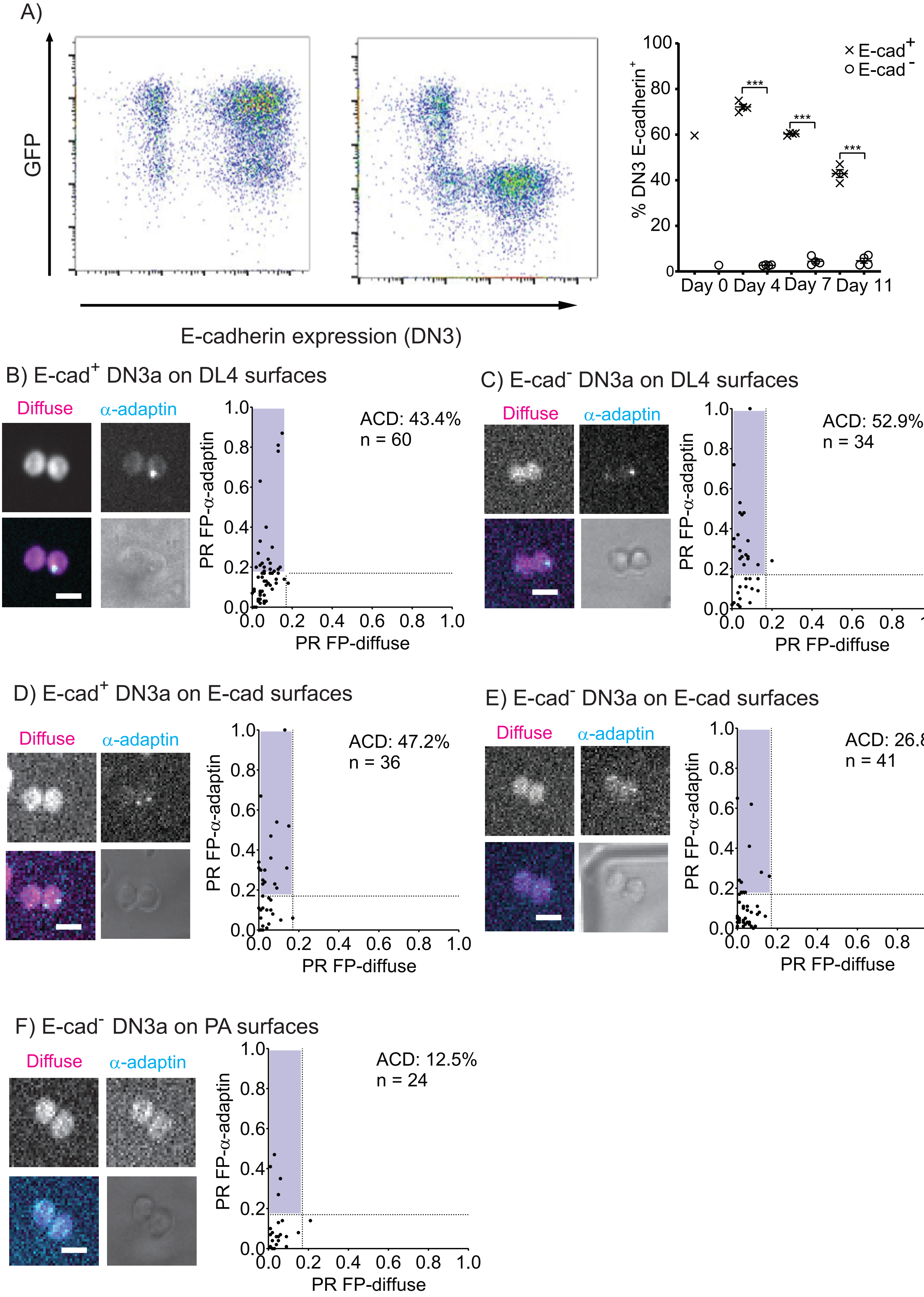

A)
GFP
E-cadherin expression (DN3)
% DN3 E-cadherin+
E-cad+
E-cad-
***
Day 0 Day 4 Day 7 Day 11
B) E-cad+ DN3a on DL4 surfaces
Diffuse
α-adaptin
ACD: 43.4%
n = 60
PR FP-α-adaptin
PR FP-diffuse
C) E-cad- DN3a on DL4 surfaces
ACD: 52.9%
n = 34
D) E-cad+ DN3a on E-cad surfaces
ACD: 47.2%
n = 36
E) E-cad- DN3a on E-cad surfaces
ACD: 26.8%
n = 41
F) E-cad- DN3a on PA surfaces
ACD: 12.5%
n = 24

Fig. S3

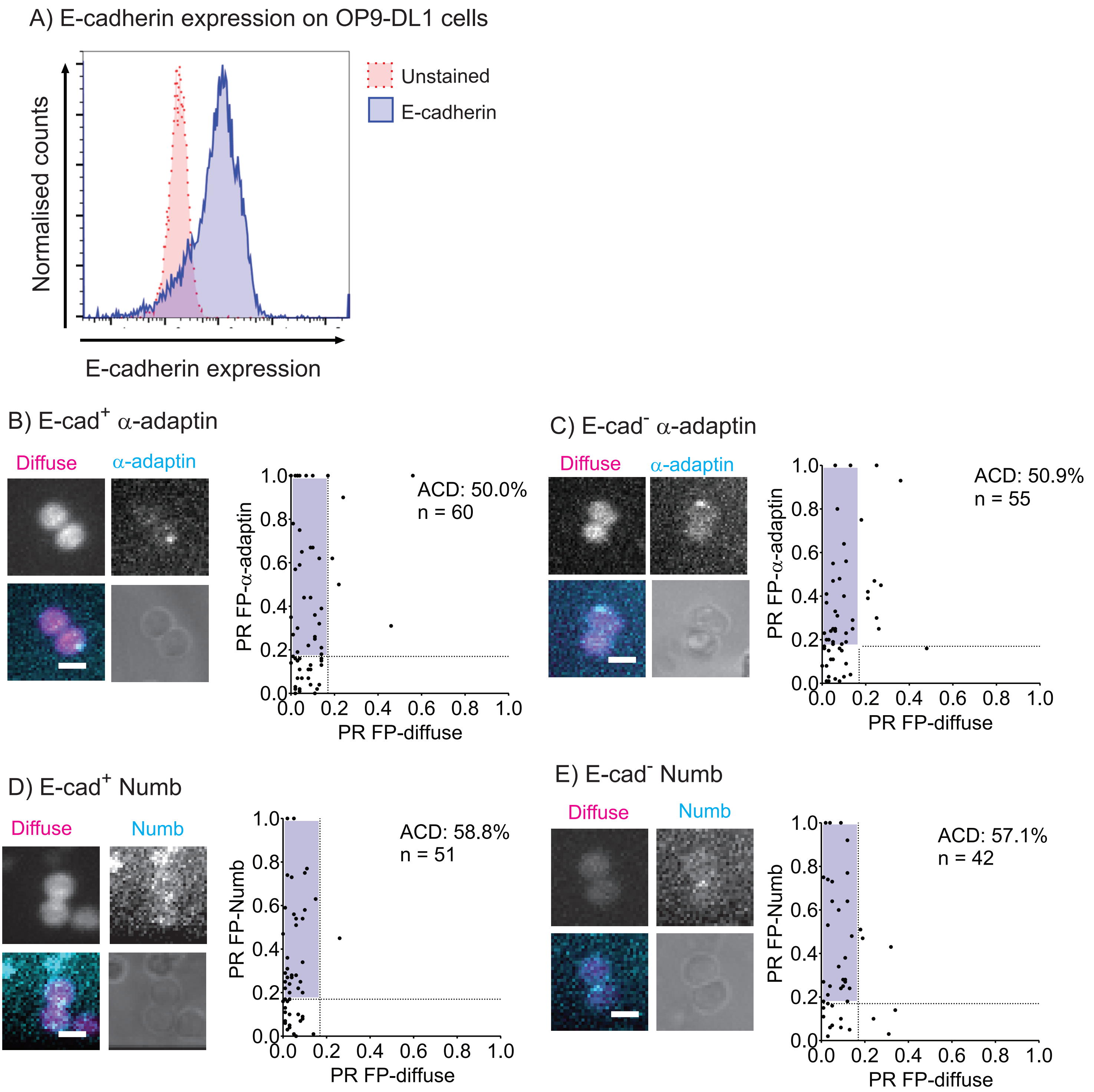

A) E-cadherin expression on OP9-DL1 cells
Unstained
E-cadherin
Normalised counts
E-cadherin expression
B) E-cad+ α-adaptin
Diffuse
α-adaptin
ACD: 50.0%
n = 60
PR FP-α-adaptin
PR FP-diffuse
C) E-cad- α-adaptin
Diffuse
α-adaptin
ACD: 50.9%
n = 55
PR FP-α-adaptin
PR FP-diffuse
D) E-cad+ Numb
Diffuse
Numb
ACD: 58.8%
n = 51
PR FP-Numb
PR FP-diffuse
E) E-cad- Numb
Diffuse
Numb
ACD: 57.1%
n = 42
PR FP-Numb
PR FP-diffuse

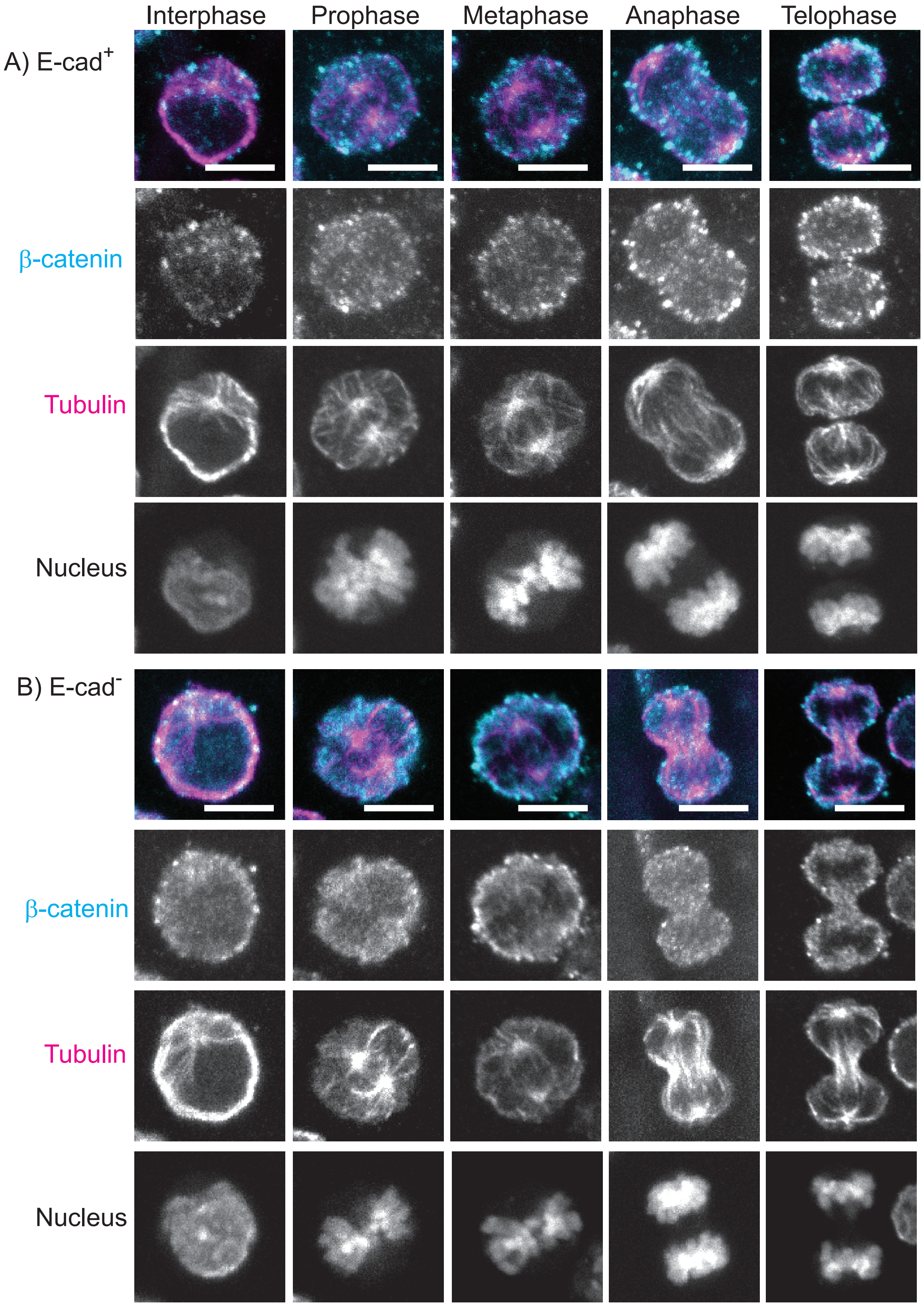
Interphase
Prophase
Metaphase
Anaphase
Telophase
A) E-cad+
β-catenin
Tubulin
Nucleus
B) E-cad-
β-catenin
Tubulin
Nucleus

Fig. S5

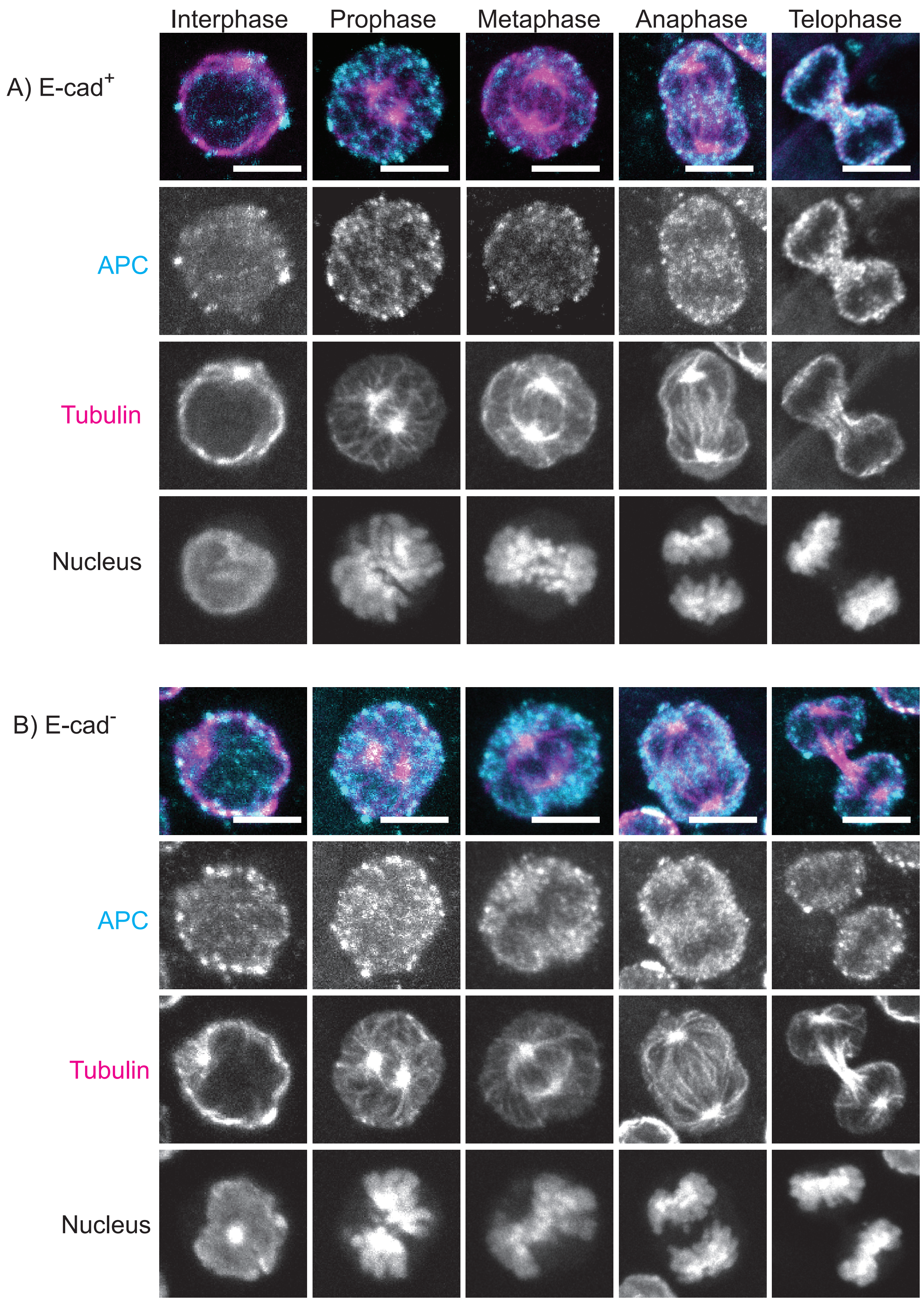

Interphase
Prophase
Metaphase
Anaphase
Telophase
A) E-cad+
APC
Tubulin
Nucleus
B) E-cad-
APC
Tubulin
Nucleus

Fig. S6

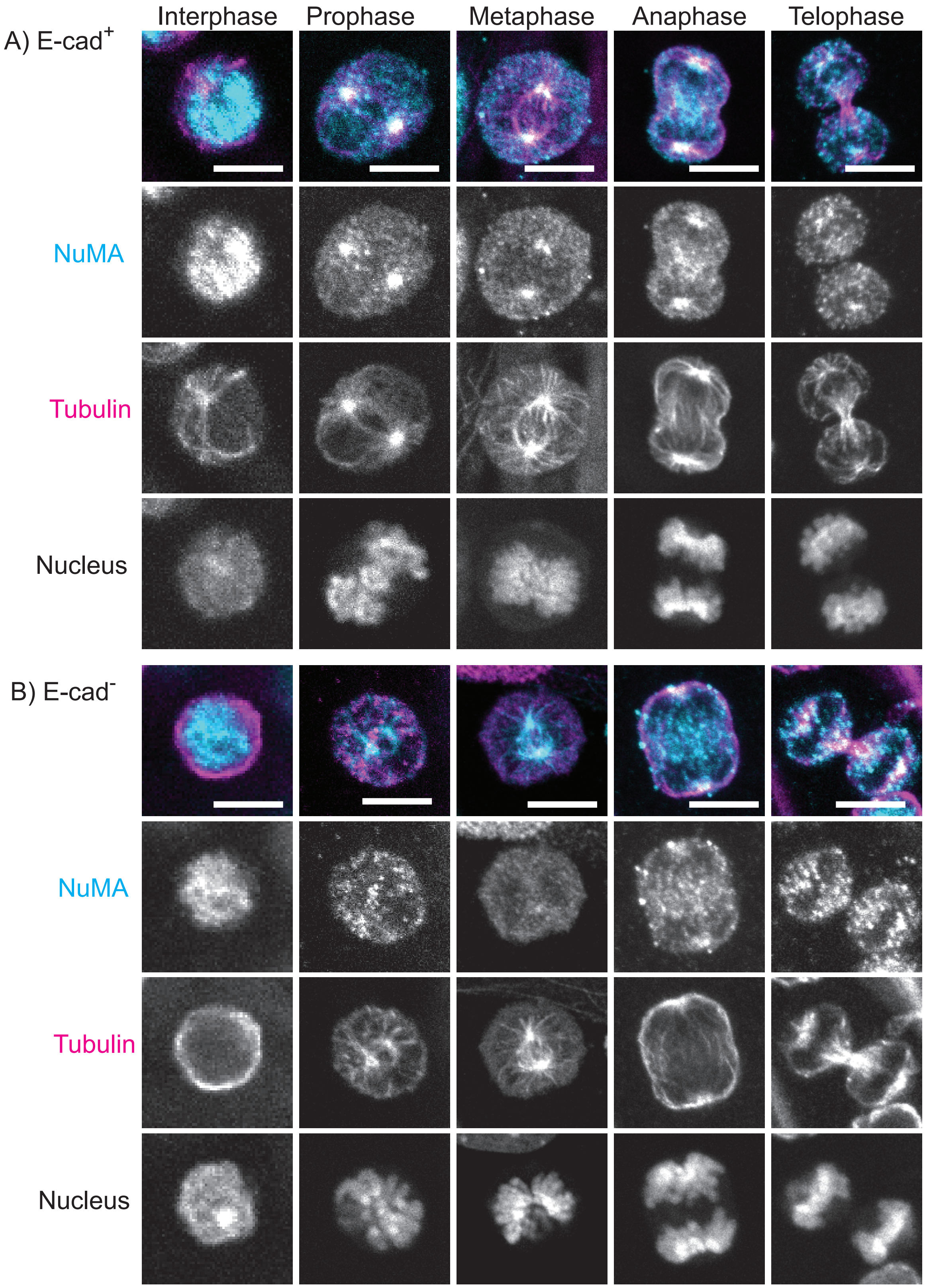

Interphase
Prophase
Metaphase
Anaphase
Telophase
A) E-cad+
NuMA
Tubulin
Nucleus
B) E-cad-
NuMA
Tubulin
Nucleus

Fig. S7

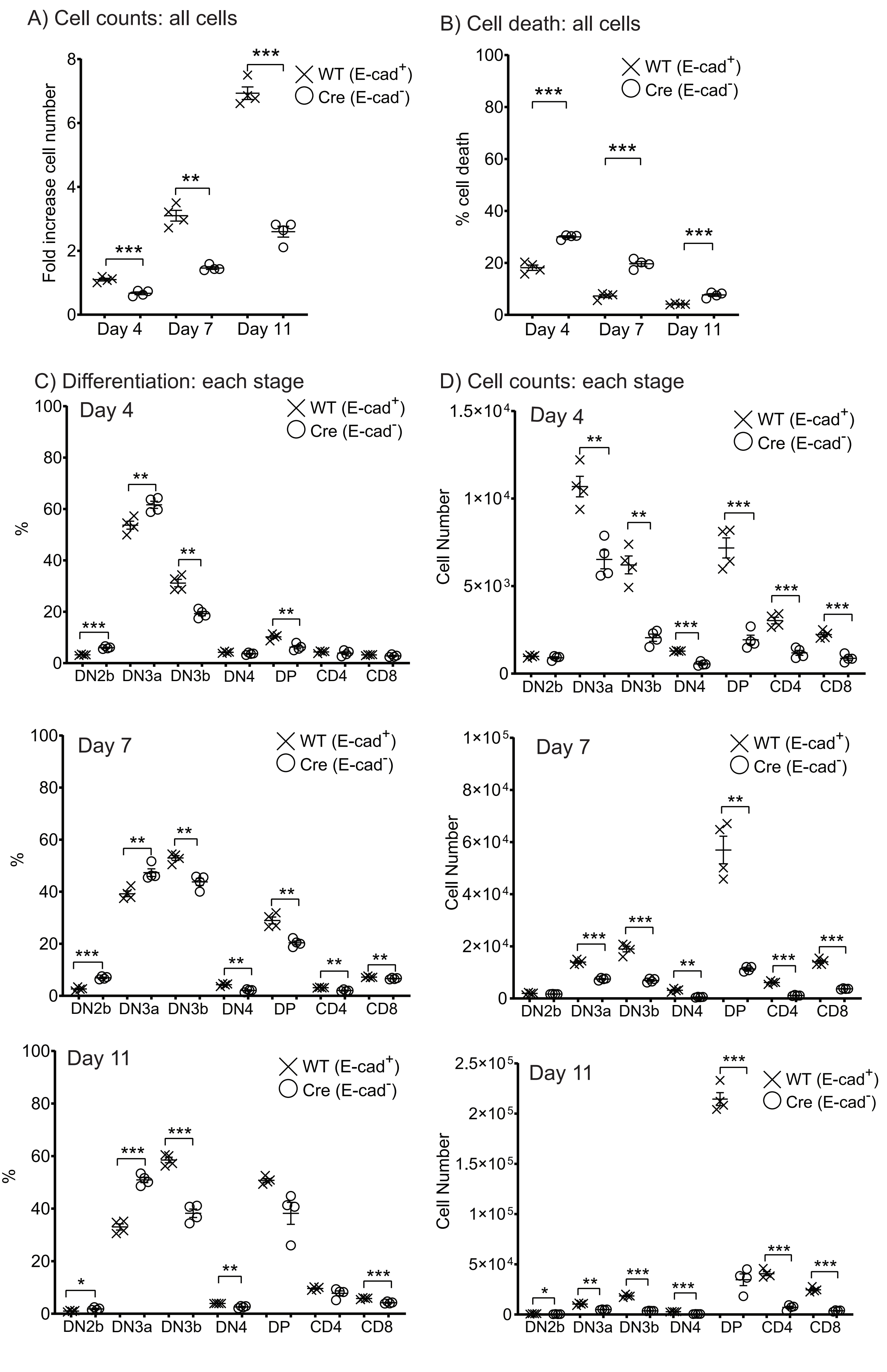
A) Cell counts: all cells
Fold increase cell number
Day 4
Day 7
Day 11
WT (E-cad+)
Cre (E-cad-)
B) Cell death: all cells
% cell death
C) Differentiation: each stage
%
DN2b
DN3a
DN3b
DN4
DP
CD4
CD8
D) Cell counts: each stage
Cell Number

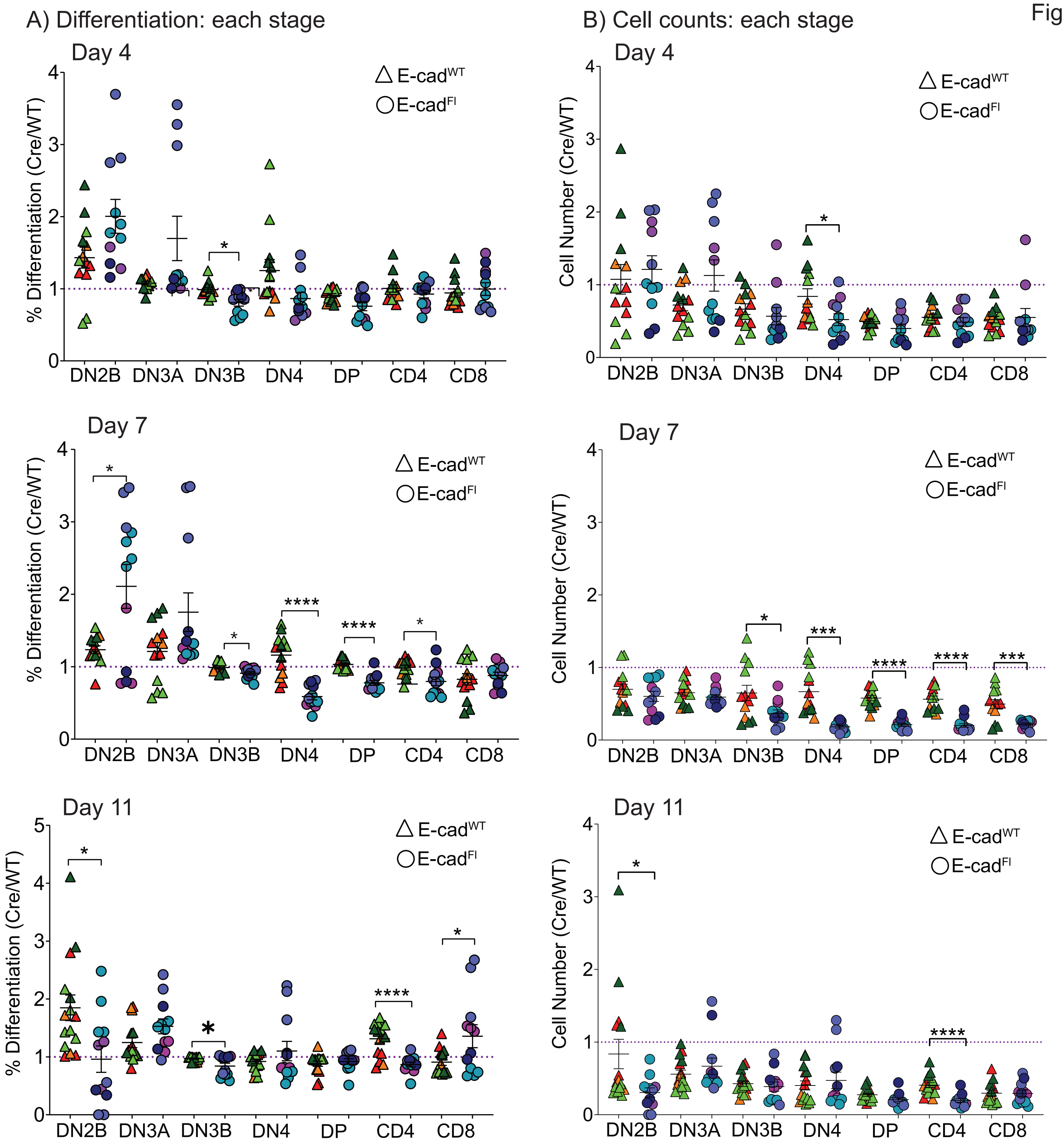
A) Differentiation: each stage
B) Cell counts: each stage
Day 4
Day 7
Day 11
% Differentiation (Cre/WT)
Cell Number (Cre/WT)
△ E-cadWT
○ E-cadFl
DN2B
DN3A
DN3B
DN4
DP
CD4
CD8
*
****
***

Fig. S9

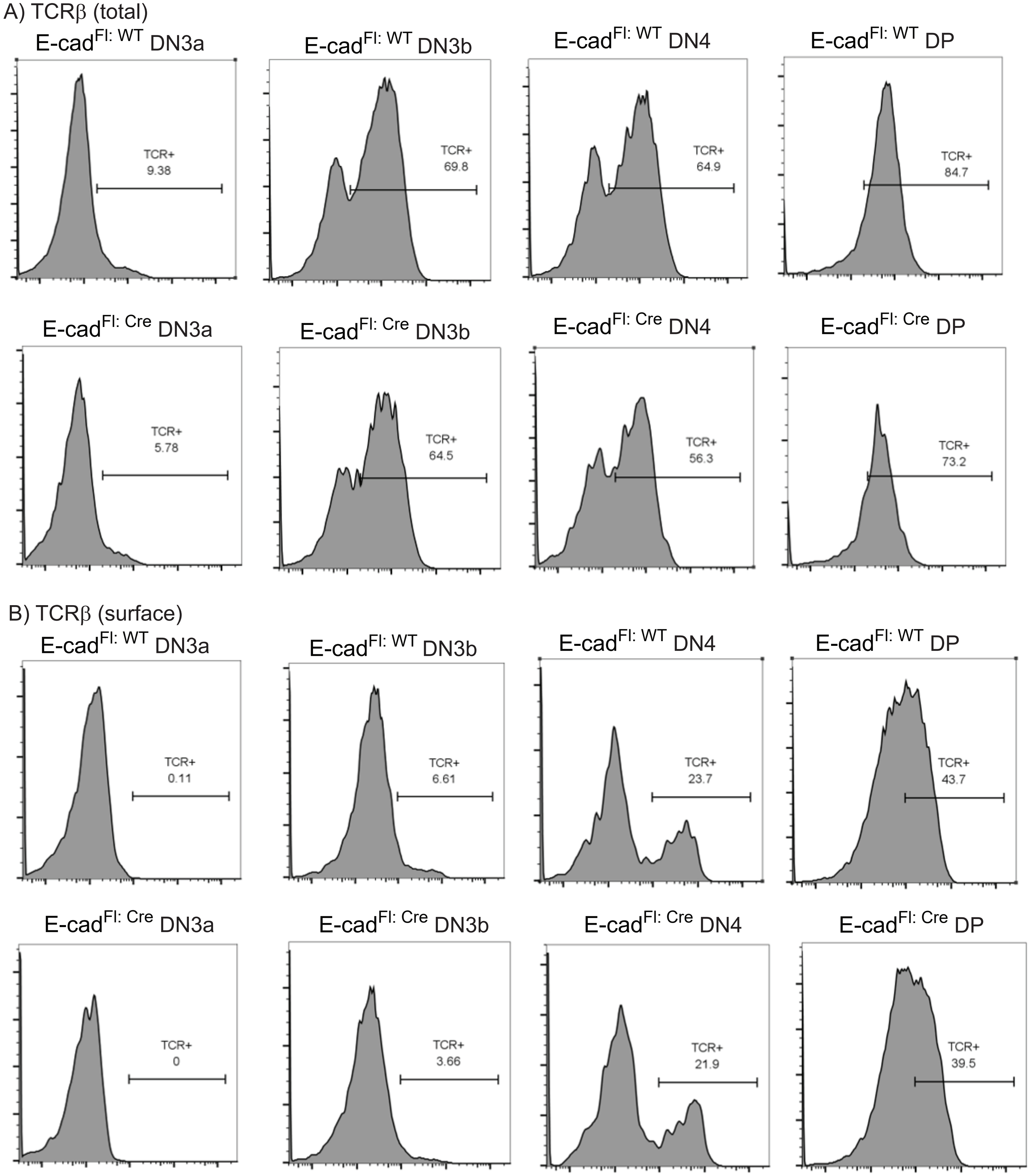
A) TCRβ (total)
E-cadFl: WT DN3a
TCR+
9.38
E-cadFl: WT DN3b
TCR+
69.8
E-cadFl: WT DN4
TCR+
64.9
E-cadFl: WT DP
TCR+
84.7
E-cadFl: Cre DN3a
TCR+
5.78
E-cadFl: Cre DN3b
TCR+
64.5
E-cadFl: Cre DN4
TCR+
56.3
E-cadFl: Cre DP
TCR+
73.2
B) TCRβ (surface)
E-cadFl: WT DN3a
TCR+
0.11
E-cadFl: WT DN3b
TCR+
6.61
E-cadFl: WT DN4
TCR+
23.7
E-cadFl: WT DP
TCR+
43.7
E-cadFl: Cre DN3a
TCR+
0
E-cadFl: Cre DN3b
TCR+
3.66
E-cadFl: Cre DN4
TCR+
21.9
E-cadFl: Cre DP
TCR+
39.5

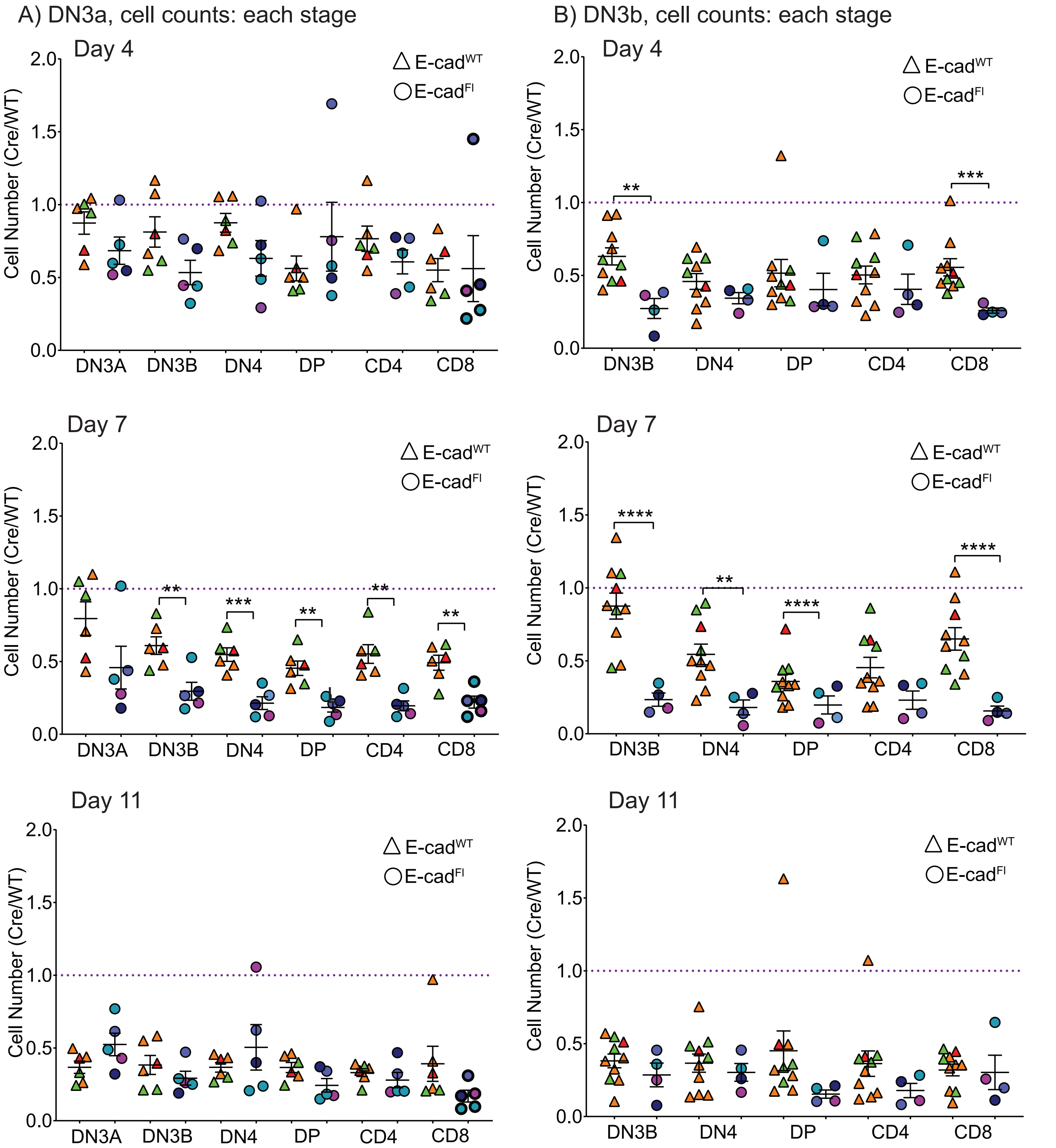
A) DN3a, cell counts: each stage
B) DN3b, cell counts: each stage
Day 4
Day 7
Day 11
E-cadWT
E-cadFl
Cell Number (Cre/WT)
2.0
1.5
1.0
0.5
0.0
DN3A
DN3B
DN4
DP
CD4
CD8
**
***
****
